# Comparative chromatin accessibility upon BDNF-induced neuronal activity delineates neuronal regulatory elements

**DOI:** 10.1101/2021.05.28.446128

**Authors:** Ignacio L. Ibarra, Vikram S. Ratnu, Lucia Gordillo, In-Young Hwang, Luca Mariani, Kathryn Weinand, Henrik M. Hammarén, Martha L. Bulyk, Mikhail M. Savitski, Judith B. Zaugg, Kyung-Min Noh

**Affiliations:** European Molecular Biology Laboratory (EMBL), Structural and Computational Biology Unit. 69117 Hedelberg, Germany; European Molecular Biology Laboratory (EMBL), Genome Biology Unit. 69117 Heidelberg, Germany; Collaboration for joint PhD degree between EMBL and Heidelberg University, Faculty of Biosciences; Institute of Computational Biology, Helmholtz Center Munich, 85764 Oberschleißheim, Germany; Division of Genetics, Department of Medicine, Brigham and Women’s Hospital and Harvard Medical School, Boston, Massachusetts 02115, USA; Department of Pathology, Brigham and Women’s Hospital and Harvard Medical School, Boston, Massachusetts 02115, USA

## Abstract

Neuronal activity induced by brain-derived neurotrophic factor (BDNF) triggers gene expression, which is crucial for neuronal survival, differentiation, synaptic plasticity, memory formation, and neurocognitive health. However, its role in chromatin regulation is unclear. Here, using temporal profiling of chromatin accessibility and transcription in mouse primary cortical neurons upon either BDNF stimulation or depolarization (KCl), we identify features that define BDNF-specific chromatin-to-gene expression programs. Enhancer activation is an early event in the regulatory control of BDNF-treated neurons, where the bZIP motif-binding Fos protein pioneered chromatin opening and cooperated with co-regulatory transcription factors (Homeobox, EGRs, and CTCF) to induce transcription. Deleting cis-regulatory sequences decreased BDNF-mediated Arc expression, a regulator of synaptic plasticity. BDNF-induced accessible regions are linked to preferential exon usage by neurodevelopmental disorder-related genes and heritability of neuronal complex traits, which were validated in human iPSC-derived neurons. Thus, we provide a comprehensive view of BDNF-mediated genome regulatory features using comparative genomic approaches to dissect mammalian neuronal activity.

## Introduction

The brain-derived neurotrophic factor (BDNF) plays a role in neuronal growth, survival, differentiation, repair, maturation, activity-induced synaptic plasticity, and memory formation (Park & Poo, 2013; Panja & Bramham, 2014; Leal *et al*, 2015). BDNF is a major source of neuronal stimulation, being synthesized and secreted in the central nervous system. BDNF-related molecular pathways are also potential targets for treating brain diseases (Choi *et al*, 2018; Nagahara *et al*, 2009) as impairments of BDNF-mediated cellular function are linked to several neurological and psychiatric disorders (Lima Giacobbo *et al*, 2019; Björkholm & Monteggia, 2016). Exogenous BDNF application in cultured cortical neurons mimics many of the *in vivo* effects of BDNF, from generating activity-dependent cellular signals to changing the spine morphology associated with long-term potentiation, thus much is known about these processes (Park & Poo, 2013; Panja & Bramham, 2014; Leal *et al*, 2015). Extracellular BDNF binds its cognate TrkB receptor (Soppet *et al*, 1991) and induces signal transduction pathways involved in phospholipase Cγ, phosphatidylinositol 3-kinase, and mitogen-activated protein kinase (MAPK), leading to the activation of transcription factors (TFs) such as the cAMP-responsive element-binding TF (CREB), early growth response factors (EGRs), and FOS transcription factor (Esvald *et al*, 2020; Minichiello, 2009; Calella *et al*, 2007). While the genomic distribution of BDNF-induced TFs is poorly characterized, collective actions of these TFs at promoters and enhancers may coordinate gene expression required for long-lasting structural and functional changes in neurons.

Chromatin regulation is a critical initial step in gene expression, however, chromatin responses to BDNF stimulation have not been analyzed. It is also unclear if the BDNF-induced chromatin changes are related to specific brain disorders and if changes in chromatin features induced by BDNF differ from other stimulatory events. Comparative analysis of epigenomic data has helped define the role of regulatory elements in various cell types (Boix *et al*, 2021), yet it has not been applied in neurons with BDNF stimulation.

For a comparative analysis of chromatin responses, neuronal activity induced by an elevated level of extracellular potassium chloride (KCl) is appealing for several reasons. While BDNF stimulation activates mainly the MAPK signaling pathway, KCl stimulation induces membrane depolarization and intracellular calcium rise (Greer & Greenberg, 2008), which triggers a series of calcium-dependent signaling events resulting in activation of TFs in the nucleus. KCl stimulation has been used to study activity-dependent gene expression and chromatin response in primary neurons (Kim *et al*, 2010; Kitazawa *et al*, 2021) and revealed crucial TFs. The TFs transcriptionally activated by KCl include neuronal PAS domain protein 4 (NPAS4), FOS, and EGRs (Yap & Greenberg, 2018), many of which overlap with BDNF stimulation.

Here, we profiled genome-scale changes in chromatin accessibility and transcription in mouse primary cortical neurons following stimulation by BDNF or KCl. Our data analyses revealed changes in chromatin accessibility and the impact of these changes on gene expression in response to BDNF, compared to depolarization by KCl. Verifying our analyses with proteomics, genome-editing, and hiPSCs-derived cortical neurons provide a clearer systemic understanding of stimulation-induced chromatin response in neurons. Our results also underscore how neuronal activity transferred to the chromatin signature connects to a functionality of human non-coding regulatory regions associated with brain disorders.

## Results

### BDNF triggers a biphasic expression response

We monitored chromatin and transcription changes across multiple time-points after BDNF stimulation in mouse primary cortical neurons and compared them to KCl depolarization (**Fig. 1a**). For KCl depolarization, we applied a concentration (55 mM) of KCl that is used to induce neuronal activity (Tyssowski *et al*, 2018). For BDNF stimulation, we applied a physiological dose (10ng/mL) previously reported in the mouse brain (Yoo *et al*, 2016). Neurons treated with KCl modestly activated phosphorylated MAPK (pMAPK), whereas different concentrations (5 to 100 ng/mL) of BDNF all strongly induced pMAPK (**Fig. 1a**).

**Figure 1.**
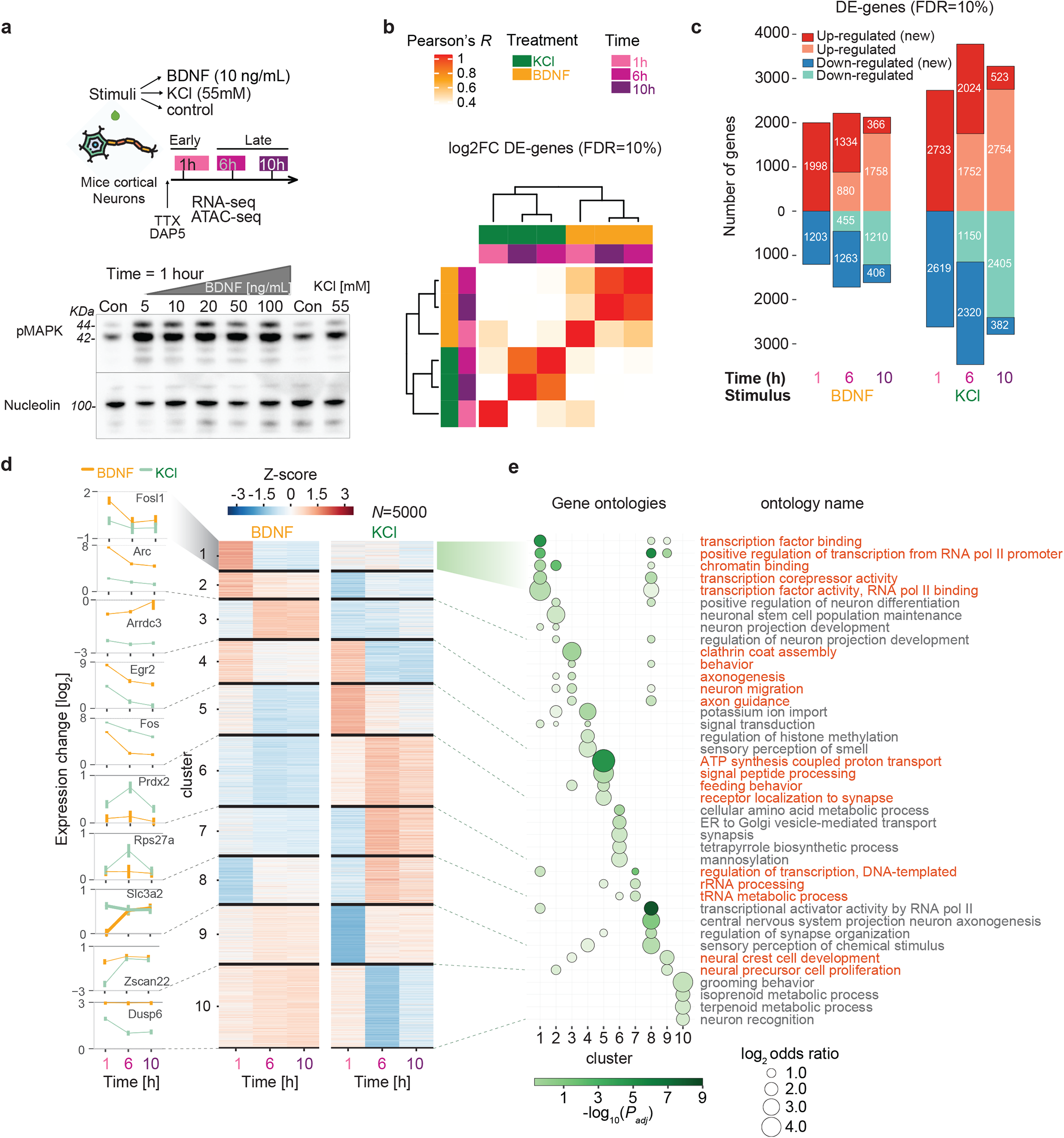
Transcriptional dynamics in mouse cortical neurons upon neuronal stimulation with BDNF and KCl defines early and late gene programs. (a) *(top)* Experimental design. Cultured cortical neurons are stimulated with BDNF, KCl or no treatment (Control) and activated neurons are prepared in three specific time points for RNA-seq and ATAC-seq at early (1h) and late time points (6 and 10h); *(bottom)* Blot for pMAPK in neurons treated with KCl 55 mM and several concentrations of BDNF (Con = no stimulation). 44 and 42 KDa bands indicated with lines. Nucleolin shown as internal control. (b) Hierarchical clustering for all differentially-expressed genes (DE-genes, adjusted P-value < 0.1, using Benjamini Hochberg’s correction) using correlation of log2-fold changes when compared to unstimulated control samples at each time point. (c) Number of DE-genes at each time-point and treatment combination (above and below zero indicates up-regulated and down-regulated genes, respectively). Darker shades indicate genes newly up/down regulated at a given time point (new). (d) Partitioning around medoids clustering-of-expression dynamics using genewise-scaled Z-scores of all DE-genes from (c) (*k* = 10 clusters). Numbers indicate clusters. Left line plots indicate expression levels for signature genes selected for each cluster. (e) Gene Ontology enrichment analysis for clusters shown in (d). Each ribbon shows clusters with their respective significant gene groups using topGO (Alexa *et al*, 2006). Up to five significant terms per cluster are shown (*P*-values adjusted with BH). Circle sizes indicate enrichment of ontology genes in each cluster versus all other clusters. Names on the right y-axis indicate ontology common names.

Using mRNA-sequencing (RNA-seq), we identified the transcriptional changes in neurons collected at 1, 6, and 10 hours after BDNF stimulation, compared to unstimulated controls (two biological replicates for each time and condition). Transcriptome profiles were highly reproducible in biological replicates (**Supplementary Figure 1a,c**). Hierarchical clustering of differentially-expressed genes (DE-genes; FDR=10%) in all conditions revealed a clear separation of BDNF-induced transcriptional changes compared to KCl-induced changes, with each stimulus inducing an early (1h) and late (6h, 10h) transcriptional response (**Fig 1b**). Comparison of the DE-genes with published RNA-seq in KCl-treated neurons (1h, 6h) (Ataman *et al*, 2016) showed a higher correlation with the KCl samples (Spearman’s ρ = 0.83/0.82 at 1/6h) than BDNF (0.76/0.21 at 1/6h) (**Supplementary Figure 1b**) validating our cell assays. Large numbers of DE-genes were observed at all times after stimulation, but DE-genes decreased at 6h and 10h in both KCl (*N =* 5,352/4,344/905 for 1, 6 and 10h, respectively) and BDNF (3,201/2,597/722) (**Fig. 1c**). Thus, while more DE-genes appeared in KCl than BDNF, both induced comparable biphasic transcriptional dynamics. The majority of DE-genes expressed at 10h were already differentially expressed at 6h, indicating stable expression of the late response genes.

Neuronal stimulation induces immediate early genes (IEGs), enriched for TFs, and delayed response genes (DRGs) associated with synaptic plasticity and neuronal function (Flavell & Greenberg, 2008; Tyssowski *et al*, 2018). To identify BDNF-induced genes, we performed unsupervised clustering of the top 5,000 significant DE-genes across stimuli and time-points, obtaining early (1h) and late (6/10h) clusters across stimuli (**Fig. 1d**). Among four clusters of early up-regulated genes, cluster 1 (e.g., Fosl1) and cluster 2 (Arc) were specific for BDNF, cluster 4 (Egr2) was increased in both stimuli, and cluster 5 (Fos) was more highly expressed in KCl. Delayed up-regulated genes specific for BDNF contained signaling-linked genes (e.g., Arrdc3, Cebpb; cluster 3) while those specific for KCl contained solute transporters and ion channel-related genes (Slc3a2, Cacna1d, Slc25a25, Kcne4, etc.; clusters 6, 7, and 8). Only a few clusters of down-regulated genes were seen in BDNF (cluster 8) and KCl (clusters 9 and 10). Gene Ontology (GO) enrichment analysis showed that clusters of early BDNF-induced genes were enriched for transcription factors (TFs), DNA-binding, and transcriptional regulation related processes as seen previously (Flavell & Greenberg, 2008; Tyssowski *et al*, 2018) (**Fig. 1e**). Delayed gene clusters showed enrichment of different GO neuronal terms. For example, axonogenesis and neuron migration appeared in BDNF (cluster 3), and many genes related to signal peptide processing, endoplasmic reticulum transport, and sensory perception were increased in KCl, but not in BDNF (clusters 6 and 8). Genes down-regulated by KCl but modestly up-regulated by BDNF (clusters 9 and 10) were enriched for neuronal cell division, neuronal recognition, metabolic process, and behavioral terms. Thus, BDNF and KCl stimulation separately trigger a biphasic transcriptional response similar in dynamics but involving different sets of genes, which could be in part mediated by variable expression of early induced TFs activating different sets of late response genes.

### BDNF alters regulatory elements within chromatin

To understand the regulatory basis of BDNF- and KCl-induced gene expression, we quantified chromatin accessibility dynamics in the same samples (**Fig. 1a**) using the assay for transposase-accessible chromatin sequencing (ATAC-seq) (Buenrostro *et al*, 2015). Principal component analysis showed that variability in chromatin peaks was reproducible within biological replicates (**Supplementary Figure 2a**). We identified a total of 58,724 peaks, of which 15,566 were differentially accessible (DA-peaks; FDR=10%) across any condition. Clustering of DA-peaks showed a distinct separation between stimuli, but unlike the DE-gene results, chromatin response upon BDNF induction was not clearly separated into early and late responses (**Fig. 2a**). We separated DA-peaks as gained DA-peaks (increased chromatin accessibility compared to control) and closing DA-peaks (decreased chromatin accessibility compared to control) and classified by the time of their first occurrence. We found that the majority of all gained DA-peaks (6,379; 88%) in BDNF appeared at 1h and more than half of them maintained increased accessibility at later time points (**Fig. 2b**). Only 446 (6.2%) and 418 (5.8%) additional peaks appeared at 6 and 10h, respectively. In contrast, the gained DA-peaks in KCl were balanced across all time points with 2,168 (46%), 1,551 (33%) and 1,025 (21%) at 1, 6 and 10h, respectively. For the closing DA-peaks, a similar pattern was seen for both BDNF and KCl: decreasing numbers from early to late time-points, and comparable fractions of newly closing DA-peaks (2,633/672/579; 1,522/640/666 for BDNF and KCl, respectively) (**Fig. 2b**). Thus, BDNF stimulation triggered rapid and extensive increases in chromatin accessibility.

**Figure 2.**
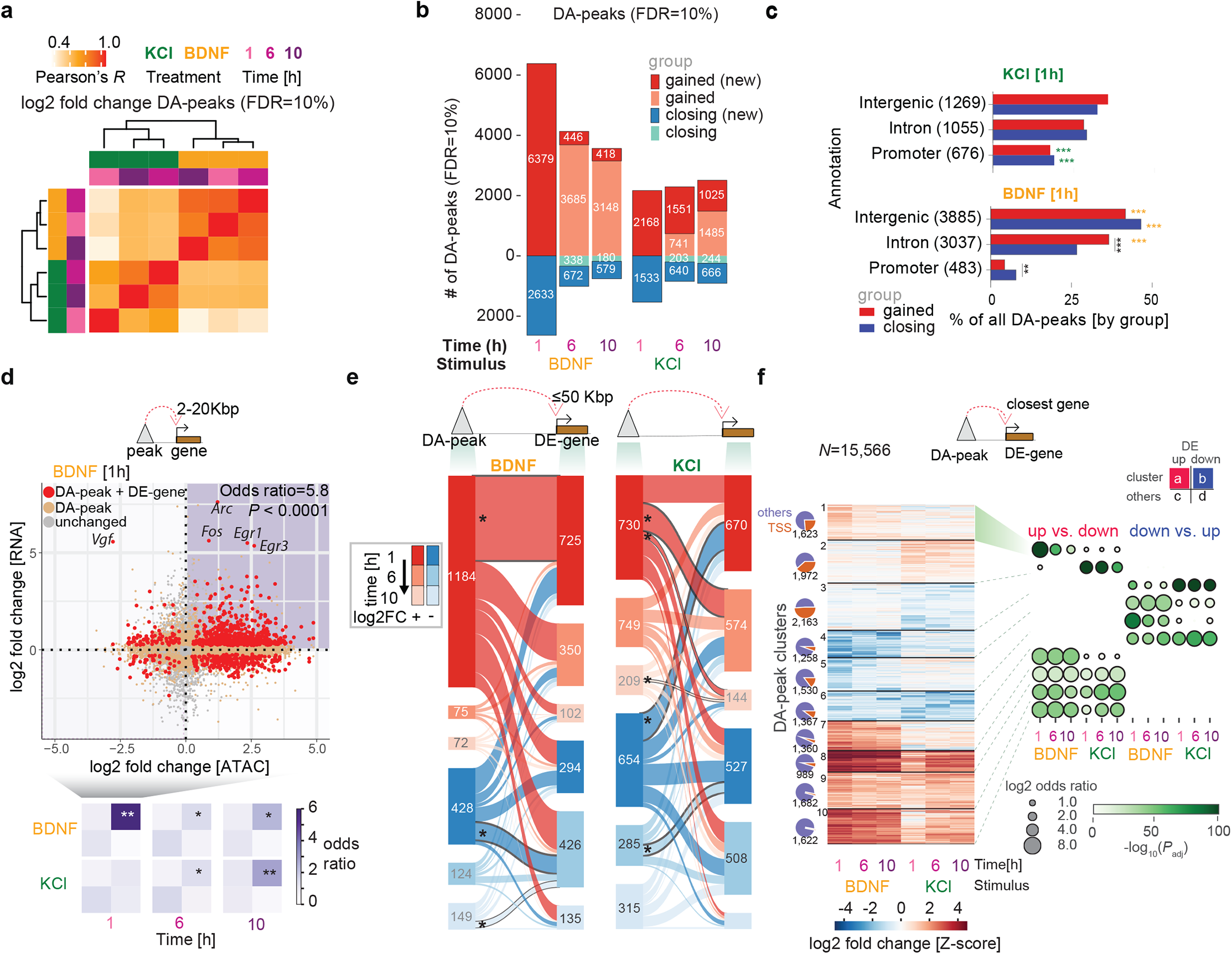
Chromatin accessibility changes upon neuronal activitaction with BDNF and KCl reveals early BDNF regulatory control of gene expression. (a) Hierarchical clustering for all differentially accessible peaks (DA-peaks, adjusted *P* < 0.1, using Benjamini Hochberg’s correction) using correlation of log2-fold changes when compared versus matched control samples. (b) Number of DE-peaks at each time-point and treatment combination (above and below zero indicates gained and closing DA-peaks, respectively). Darker shades indicate peaks newly gaind/closing at a given time point (new). (c) Percentage of gained and closing DA-peaks grouped by top genomic annotations at the 1h time-point (by percentage). Numbers in parenthesis next to each annotation label indicate the absolute number of peaks associated with the annotation. Gray asterisks indicate within-stimulation Fisher’s exact test comparisons. Green/yellow asterisks indicate between-stimulation comparisons, wherever one peak set is significantly enriched for one annotation (\*\**P* < 0.01; \*\*\**P* < 0.001). (d) (*top*) Association between distal regulatory elements (DREs) and gene expression at the BDNF 1h time-point. Each point indicates the log2-fold change of an ATAC-seq peak (x-axis) and the gene expression of a closest gene (y-axis) with distance between 2-20 Kbp. Colors indicate whether none (gray), only the peak (orange), or both peak and gene (red) show significant changes versus control neurons. (*bottom*) Enrichment for paired DA-peak and DE-gene in the four quadrants are summarized for BDNF and KCl. Asterisks indicate *P* values as corrected by a Benjamini Hochberg procedure (* = *P* < 0.05; ** = *P* < 0.01). (e) Number of associations between DA-peaks and DE-genes for KCl (left) and BDNF (right). Sankey plot shows closest DE-genes with distances <50 Kbp to DA-peaks. Asterisks indicate associations between DA-peaks and DE-genes with Z-scores >2.5 for both genes and peaks, using a permutation approach for DA-peaks and their connected DE-genes while maintaining timewise changes (see Methods). (f) (*left*) Partitioning around medoids clustering of accessibility dynamics using scaled Z-scores for log2 fold changes of all DA-peaks in (b)(*k* = 10 clusters). Venn diagrams indicate proportion of peaks in each cluster associated to TSS or any other regulatory region (*right*) Enrichment of up-regulated (DE-up) versus down-regulated (DE-down) DE-genes (up vs. down), and enrichment of DE-down versus DE-up genes (down vs. up) in closest genes within a cluster when compared with the same category in other clusters. Enrichments calculated using Fisher’s exact test, with adjusted P-value correction via Benjamini Hochberg).

Despite the higher number of DE-genes at 1h in KCl, we found over two-fold more DA-peaks at 1h in BDNF (9,012) relative to KCl (3,701) (**Fig. 2b; Supplementary Figure 2b**). These results imply that BDNF stimulation may induce a regulatory remodeling of the chromatin landscape, whereas KCl has a more substantial effect on transcription. Indeed, when DA-peaks at 1h were subdivided into intergenic, intronic, and gene promoter regions, DA-peaks in KCl were over-represented in promoters (20% for KCl and 5% for BDNF; Fisher’s exact test adjusted *P* value < 0.001; two-sided) and BDNF peaks were enriched for intergenic regions and introns (32% for KCl and 47% for BDNF; adjusted *P* < 0.001) (**Fig 2c**, all annotations in **Supplementary Figure 2c**). Chromatin state modeling (ChromHMM (Ernst & Kellis, 2012)) analysis based on neuronal datasets (Su *et al*, 2017) further revealed that gained DA-peaks in BDNF were associated with enhancers (over 10-fold more enrichment than KCl at genic enhancers). Gained DA-peaks in KCl were located at active transcription start sites (TSS, highest log2 fold enrichment 2.3 in KCl vs. 1.4 in BDNF) and bivalent promoters marking neuronal genes (log2 fold enrichment 1.1 vs 0.1) (**Supplementary Figure 2d**). Fewer differences were observed in the closing DA-peaks except for CCCTC-binding factor (CTCF)-associated regions, which was significantly enriched in KCl (log2 fold enrichment 2.0, 2.1-fold higher than BDNF). Thus, the two responses promote different chromatin regulatory architectures early on, with KCl affecting gene promoters and BDNF acting preferentially through regulatory elements, such as enhancers.

Based on a study linking neuronal activity response to the three-dimensional conformation of the genome (Beagan *et al*, 2020), we integrated the DA-peaks with published Hi-C maps generated across neurodevelopment (Bonev *et al*, 2017). We observed higher correlations for loop-associated peak pairs in cortical neurons (**Supplementary Figure 3a**), suggesting a role for genome topology in co-varying regions. Furthermore, co-variation of enhancer accessibility and gene expression is enriched for HiC contacts at 1h in BDNF. An equivalent analysis for KCl showed a similar trend which was, however, not statistically significant (**Supplementary Figure 3b**). Thus, following BDNF stimulation, changes in enhancer accessibility appear to translate into gene expression changes and these correlate with changes in the physical connectivity in the genome.

### Neuronal chromatin dynamics affect gene expression

We further investigated the functional relationship between chromatin accessibility changes at regulatory elements and transcription, observing a significant association at specific time points between gained accessibility and increased gene expression upon BDNF (2-20 Kbp from TSS, enhancers in **Fig. 2d;** 0-2 Kbp from TSS, promoters in **Supplementary Figure 4a-b**). The strongest association was between gained DA-peaks and up-regulated DE-genes at 1h after BDNF induction (odds ratio = 5.8 for enhancers; 20.8 for promoters; adjusted P = 2 x 10^-18^, two-sided Fisher’s exact test with BH correction). Weaker associations were observed in enhancers at 6h and 10h after BDNF (OR = 1.7 at 6h; 2.3 at 10h; **Fig. 2d**, bottom). KCl showed no significant association between gained DA-peaks and DE-genes in promoters or enhancers at 1h, but gained DA-peaks and up-regulated DE-genes became increasingly associated in enhancers at later time points (OR = 1.6 at 6h; 2.5 at 10h; P < 0.01). This suggests that gained accessibility in promoters is linked to up-regulation of genes at 1h after BDNF, whereas for KCl, the gene expression outcome of gained DA-peaks in promoters is more complex. Chromatin remodeling at distal regulatory elements has a rapid impact on BDNF-induced gene expression, while for KCl, it affects gene up-regulation at 6h and 10h. No significant association was observed for closing peaks and down-regulated genes at each time points.

To examine whether early changes in accessibility prime gene expression at later consecutive time points, we analyzed the association between DA-peaks (1h, 6h, 10h; gained new and closing new in **Fig. 2b**) and their nearest DE-gene (50 Kbp or less), classified peak-to-gene associations by the time of their first appearance, and assessed whether any pair of peak-to-gene classes were over-represented (using Z-scores based on peak label permutations; see **Methods**; **Fig. 2e**). We found that gained DA-peaks at 1h post-BDNF induction were significantly correlated only to genes already up-regulated at 1h (Z-score > 6; **Supplementary Figure 4c; Supplementary Data 2**) without being correlated with newly induced DE-genes at later times, whereas decreasing DA-peaks at 1h BDNF were correlated to newly down-regulated genes at 6h (Z-score > 3). For KCl, in contrast, early increased DA peaks (1h) were significantly associated with newly induced DE genes at 6h and 10h (Z-scores > 2.5). Thus, increased and decreased chromatin accessibilities at 1h likely prime gene expression at later times in KCl and BDNF, respectively.

To further dissect the relationship between chromatin accessibility and gene expression we performed unsupervised clustering of all DA-peaks across time points and conditions (KCl and BDNF), grouped them into 10 clusters (**Fig. 2f**) and calculated whether they were enriched for peaks connected to up- versus down-regulated genes at individual time points and conditions. A set of BDNF-specific early-response peaks associated with early-response genes in BDNF (cluster 1; 1,623 peaks) and a similar peak cluster for KCl (cluster 2; 1,972 peaks). Both cluster 1 and 2 contain a large fraction of promoters. Sets of shared DA-peaks that were affected by both BDNF and KCl (clusters 8-10; 4,293 peaks) showed a faster response in BDNF than in KCl for accessibility and gene expression, and are comprised mainly of distal elements. Taken together, BDNF induces robust changes in chromatin accessibility that directs early gene expression, which persists to later time-points. In contrast, accessibility changes appear delayed in the KCl response, and specific chromatin patterns were only enriched for late-response genes.

### TF motifs underlying chromatin responses

Given that chromatin accessibility in distal elements was partially shared between BDNF and KCl (despite a delay in the KCl response), we explored common and specific TF activity after stimulation. To identify TF binding motifs in each set of gained and closing DA-peaks, we used 8-mers describing 108 TF specificity groups (Mariani *et al*, 2017) and a position weight matrices (PWMs) database for TF binding specificities (Weirauch *et al*, 2013), quantifying TF motif enrichment in comparison with mouse-specific negative control regions (generated by GENRE (Mariani *et al*, 2017); **Fig. 3a**; **Methods**). Some 69% of DA-peaks appearing after stimulation are explained by one of the 16 over-represented TF motifs (**Supplementary Figure 5a**) indicating a small set of TFs dominate the activity-dependent regulatory landscape. The basic leucine zipper (bZIP) domain motif showed the highest motif enrichment in gained DA-peaks for both BDNF and KCl (**Fig. 3b**, receiver operating characteristic area under the curve ([ROC-AUC] = 0.65; P < 0.0001; Wilcoxon rank sum test, BH-adjusted), consistent with an *in vivo* study of electroconvulsive stimulation in mouse brain (Su *et al*, 2017). The prominent effect of bZIP on opening chromatin regions was corroborated by the physical centrality of bZIP sites in gained DA-peaks (**Supplementary Figure 5b**) consistent with a pioneer role for these TFs (Vierbuchen *et al*, 2017; Su *et al*, 2017). Homeobox (Hbox-III) and POU domain (POU; POU-HMG) motifs were also enriched in gained DA-peaks after BDNF and KCl stimulation (ROC-AUC > 0.55). Among the BDNF-gained peaks, we found motifs for the two Homeobox subgroups (Hbox and Hbox-II), early growth response (EGR), ETS, and TALE/zf-C2H2, which are mainly activator TFs. In contrast, E2F/zf-C2H2 and KLF motifs, which function both as activator and repressor TFs, were enriched at 1h post-induction in KCl only, implying a more complex gene expression outcome of gained DA-peaks at 1h upon KCl stimulation. Furthermore, closing DA-peaks were significantly associated with motifs for hypermethylated in cancer 1 (HIC1) and regulatory factor X (RFX) in BDNF, and with motifs for CTCF, E2F, KLF, and zf-CXXC/SAND in KCl (**Fig. 3b**) implying a stimulus-specific role for these TFs.

**Figure 3.**
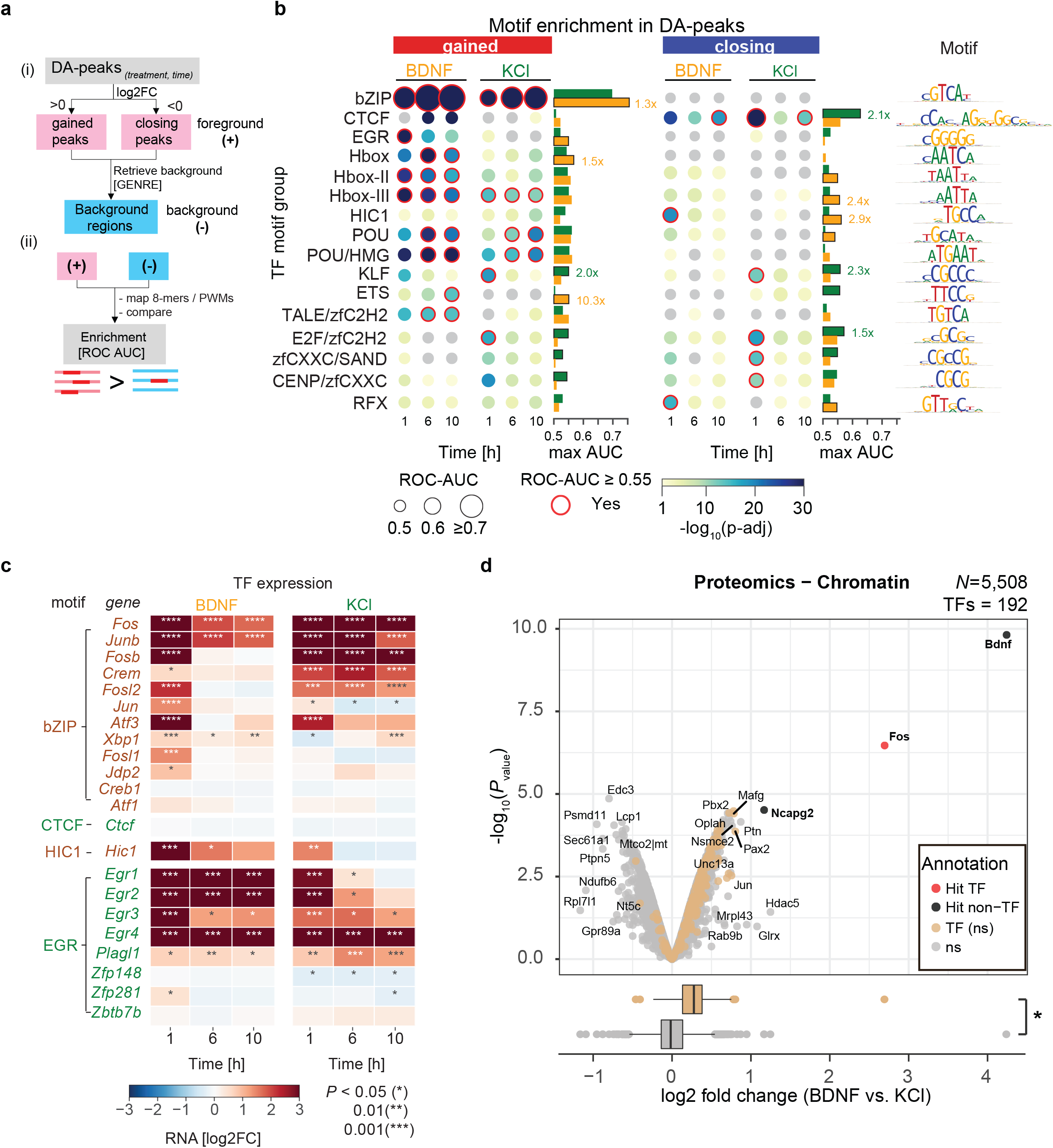
TFs linked to differentially accessible chromatin regions identify common and stimulus-specific regulatory TFs. (a) (i) DA-peaks obtained at each time point are analyzed with GENRE (Genomically Equivalent Negative REgions) software to generate regions used as negative controls in enrichment tests(Mariani *et al*, 2017). (ii) A library of 8-mers and position weight matrices (both motifs) are used to classify positive sequences (DA-peaks) versus negative regions as a receiver operating characteristic area under the curve value (ROC-AUC). (b) Enrichment of regulatory motifs in gained (*left*) and closing (*right*) differentially accessible ATAC-seq peaks. Circle size indicates ROC-AUC for recovery of positive versus negative regions. Circle color indicates *P-*value significance (one-sided Wilcoxon rank sums test). Red lines indicate adjusted *P* < 0.1 after Benjamini Hochberg’s correction. ROC-AUC values lower than 0.5 are shown as gray circles. Barplots indicate maximum value observed for each TF group across all time points for BDNF (orange bar) and KCl (green bar). In relevant cases for main text fold changes are labeled. (c) RNA expression changes for TFs related to 8-mer groups bZIP, CTCF, HIC1, and EGR. Significant changes relative to control samples are shown with asterisks. (d) Proteomics of the chromatin-bound fraction from mESC-derived neurons 1h after stimulation show significant enrichment of TFs upon stimulation with BDNF compared to KCl (boxplot, lower panel, wilcoxon two-tailed rank sum test, *P* < 0.0001). Out of 5,508 detected proteins, Fos shows a significant abundance increase after BDNF stimulation compared to KCl (|log2 fold change| ≥ 1, adjusted *P-*value < 0.05).

Given TFs of the same family share DNA-binding domains and recognize similar motifs (Mariani *et al*, 2017; Weirauch *et al*, 2014), we used RNA-seq expression of individual TFs to refine the observed motif enrichments. Among twelve TF members bound to bZIP motifs, *Fos*, *Junb*, *Fosb*, and *Fosl2* transcripts increased significantly at 1 h in both BDNF and KCl, consistent with the enrichments in gained peaks (**Fig. 3c**). Their expression levels were reduced at 6h in BDNF, but sustained in KCl until 10h, possibly reflecting the delay in activation for KCl stimulation. Among the eight TF members with EGR specificities, four (*Egr-1/2/3/4*) were induced in both BDNF and KCl at 1 h, with increased *Egr1* and *Egr2* levels continuing until 10h in BDNF but declined at 6h in KCl. The higher *Hic1* expression levels in BDNF relative to KCl, together with the *Hic1* motif being enriched in early closing DA-peaks upon BDNF stimulation, is consistent with *Hic1* acting as repressor (Pinte *et al*, 2004; Ubaid Ullah *et al*, 2018; Boulay *et al*, 2012). Despite high enrichment of the CTCF motif in KCl-induced closing DA-peaks, *CTCF* showed invariable expression levels, consistent with its ubiquitous expression and structural role in the genome (Phillips & Corces, 2009).

To assess TF protein levels, we performed mass spectrometry-based quantitative proteomic analyses on the chromatin-bound fraction at 1h post-stimulation (**Methods**). A significant increase in Fos protein abundance was observed in samples after BDNF stimulation, which was more pronounced than that in KCl (adjusted *P* < 0.2; two-sided Wald test, BH-adjusted) (**Fig. 3d, upper panel**). Fos was the highest-enriched endogenous protein among the chromatin-bound fraction after 1h BDNF stimulation. Among all 5,508 proteins detected, we observed an overall increase in the abundance of 192 combined TF proteins after BDNF stimulation compared to KCl (**Fig. 3d, lower panel**). However, the levels of the individual TFs, other than Fos, did not significantly change, possibly due to their low protein abundances. These results are consistent with the Fos protein playing a central role in neuronal activity related to synaptic plasticity, memory, and learning (Kandel, 2012). Our transcription and proteomics results are concordant with the bZIP motif being the most enriched in gained DA-peaks, and support the functional role of bZIP, especially Fos protein in augmenting chromatin accessibility upon BDNF stimulation. The increases in Fos protein were lower after 1h following KCl induction, which may contribute to the delayed response observed at enhancers in KCl.

### bZIP and TF cooperativity in induced gene expression

Chromatin regions that open early in response to bZIP could potentially be amenable to other cofactors. We tested whether the co-presence of bZIP with other TF motifs (**Fig. 3b**) increased accessibility over the bZIP motif alone (**Fig. 4a;** see **Methods**). POU, Hbox, ETS, or RFX motifs are co-present with bZIP (bZIP+TF2) and this is associated with a significant increase in accessibility after BDNF and KCl stimulation. In contrast, EGR or KLF motifs co-present with bZIP did not show any further changes in accessibility (**Fig. 4b, Supplementary Figure 6a**). To evaluate the collaborative effects of the co-present sites on gene expression, we compared the change in expression for genes near peaks with bZIP+TF2 versus peaks with bZIP alone. The co-presence of TFs such as ETS, RFX, EGR, KLF, zfCXXC, and CTCF with bZIP at proximal regions (< 2kb from TSS) of target genes was positively associated with BDNF-induced gene expression (**Fig. 4b**). EGR and KLF were associated with increased expression of proximal genes upon BDNF stimulation without any additive effect on chromatin accessibility compared to bZIP alone (**Fig. 4b**). Thus, while bZIP is the main TF responsible chromatin accessibility, other TFs, such as POU and Hbox, further enhance accessibility without directly affecting nearby gene expression, whereas EGR and KLF may be required to fine-tune BDNF induced gene expression without changing accessibility.

**Figure 4.**
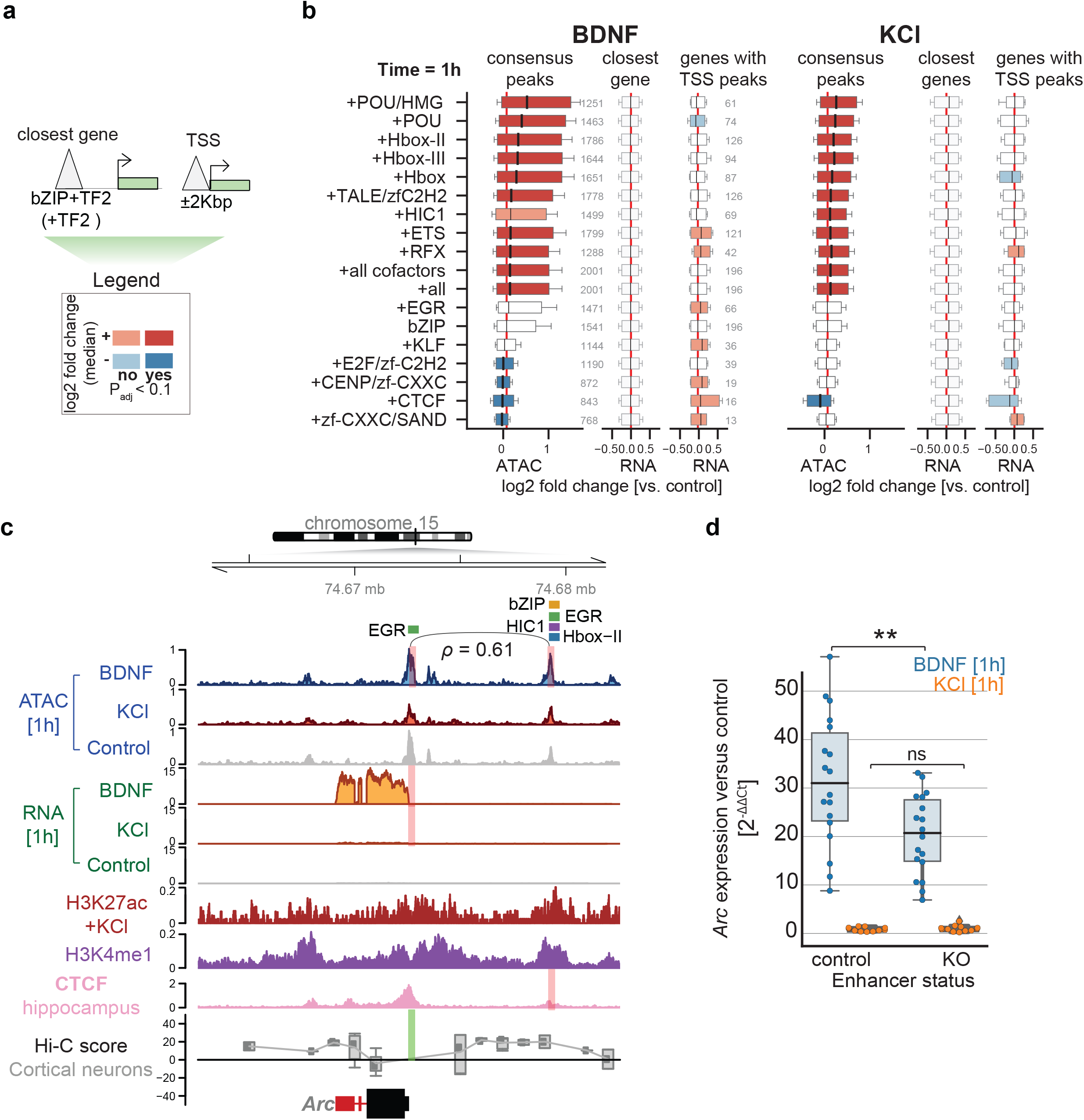
BDNF transcriptional up-regulation is linked to bZIP and EGR interactions in promoters and enhancers. (a) Scheme indicating annotation of ATAC-seq peaks based on the presence of a bZIP motif (bZIP) and another TF (TF2). Peaks are linked to genes based on closest genomic distance (“closest gene”) or only if present in gene TSS (“TSS”). A log2-fold change median indicates the difference in accessibility or expression log2-fold changes between all peaks with bZIP+TF2 motifs versus only bZIP peaks, and equivalently for the expression values of genes connecting to those peaks. (b) Genome-wide changes at bZIP+TF2 and bZIP measured as log2-fold change distributions for accessibility (“consensus peaks”), genome-wide closest genes (“closest gene”) and proximal target gene RNA levels (“genes with TSS peaks”) when bZIP motifs are co-present with other TFs (TF2) in ATAC-seq peaks versus bZIP alone. Significance is assessed using one-sided Wilcoxon rank sums test between peaks with *k*-mers co-present versus peaks with only bZIP. Absolute changes lower than 0.1 and not significant are shown as white boxplot bars. (c) Chromosome 15 genome tracks neighbouring *Arc*, displaying ATAC-seq read counts per million (CPM); RNA-seq CPM; H3K27ac normalized signal upon KCl stimulation, H3K4me1 (Malik *et al*, 2014); CTCF (Sams *et al*, 2016); and Cortical neurons Hi-C data (Bonev *et al*, 2017). Red bars in ATAC-seq tracks indicate gained DA-peaks in BDNF, and red bars in RNA-seq tracks indicate *Arc* differential expression in BDNF and KCl 1h. Green blocks in Hi-C tracks indicate anchor points for calculation of contact scores, using shaman (Cohen *et al*, 2017). Line and Spearman’s rho value indicate counts rank-based correlation between highlighted peaks across all samples (**Supplementary Figure 4c**). TF module names indicate the presence of *8*-mers in those peaks (also **Supplementary Figure 12**). (d) qPCR-based measurements of Arc expression changes in neurons treated with BDNF (blue) and KCl (orange) for control neurons, and CRISPR-Cas KOs of the proximal enhancer highlighted in **c** (N=6, 2 independent biological replicates of 3 different clonal lines).

bZIP+TF2 co-presence at distal elements (>2kb from TSS) did not show any significant effects on expression of the closest gene, which may partially be due to the complexity of mapping distal regulatory sites to their correct target genes. Cooperation was seen between a subset of increased DA-peaks at enhancers and up-regulated DA-genes (**Fig. 2d**), where co-occurrence of bZIP+TF2 could have an impact. We focused on the *Arc* gene, a key effector for synaptic function (Tzingounis & Nicoll, 2006; Plath *et al*, 2006) which is substantially induced by BDNF stimulation (**Fig. 4c**). The distal region of *Arc,* showed increased accessibility at 1h in BDNF and contains various activator TF motifs (bZIP, EGR, Hbox-II) and one repressor TF motif (HIC1) in close proximity. This distal region exhibited properties of an active enhancer (enriched H3K27ac and H3K4me1 marks obtained from neuronal epigenomics data) (Malik *et al*, 2014), bound CTCF (Sams *et al*, 2016; Ren *et al*, 2017) and had a Hi-C contact (Bonev *et al*, 2017) with the *Arc* gene promoter. Thus, we hypothesized that TF motifs adjacent to bZIP in this enhancer region could contribute to higher Arc expression upon BDNF stimulation. To assess this, we generated mouse embryonic stem cell (mESC) clones that homozygously removed the distal genomic region containing one of the three EGR motifs and one or two HIC1 motifs adjacent to the EGR motif, depending on the clone, without disturbing the other motifs (Jinek *et al*, 2012) (**Methods, Supplementary Figure 6b**). We differentiated the clones into neurons and measured *Arc* expression, using RT-qPCR, after stimulation. Similar to primary neurons, *Arc* expression increased upon BDNF stimulation in ESC-derived neurons. We observed a significant reduction in *Arc* gene expression upon BDNF stimulation in clones with a deletion of the distal TF motifs (*t*-stat = −3.0, *P* < 0.01; two-sided *t-*test), but not in CRISPR controls (*P* > 0.05) (**Fig. 4d; Supplementary Data 3**). Deleting different numbers of HIC1 motifs did not affect the level of *Arc* gene reduction upon BDNF stimulation, nor did deletion of these TF motifs show any effect on *Arc* gene expression upon KCl stimulation (**Fig. 4d**). These results suggest that an EGR motif close to bZIP in the distal regulatory element functions in BDNF-mediated *Arc* gene activation.

### Differential neuronal gene exon usage

Our TF cooperativity analysis revealed that CTCF motifs were significantly co-localized with EGR in gained DA-peaks after BDNF and KCl stimulation, and in closing DA-peaks upon BDNF stimulation (**Fig. 5a**). Most DA-peaks contain non-overlapping motifs for CTCF and EGR, suggesting the colocalization is not an artifact based on partial EGR and CTCF motif overlap. Closing DA-peaks at 1h in KCl were enriched for CTCF binding sites without EGR motifs, convergent CTCF motifs between proximal peaks in promoter-exon pairs, and known CTCF promoter-exon loop annotations (OR = 3.5; Fisher’s exact test adjusted *P* < 0.001; **Fig. 5b**). Thus, there may be a functional interaction between CTCF and EGR with looping-associated gene regulation, which might not act in closing DA-peaks upon KCl stimulation. Given that CTCF binding may affect exon usage (Shukla *et al*, 2011; Paredes *et al*, 2013; Ruiz-Velasco *et al*, 2017), these insights combined with the enrichment of CTCF binding sites in both intronic and exonic regions of DA-peaks (**Supplementary Figure 7a**), suggest activity-dependent chromatin accessibility may play a role in alternative splicing.

**Figure 5.**
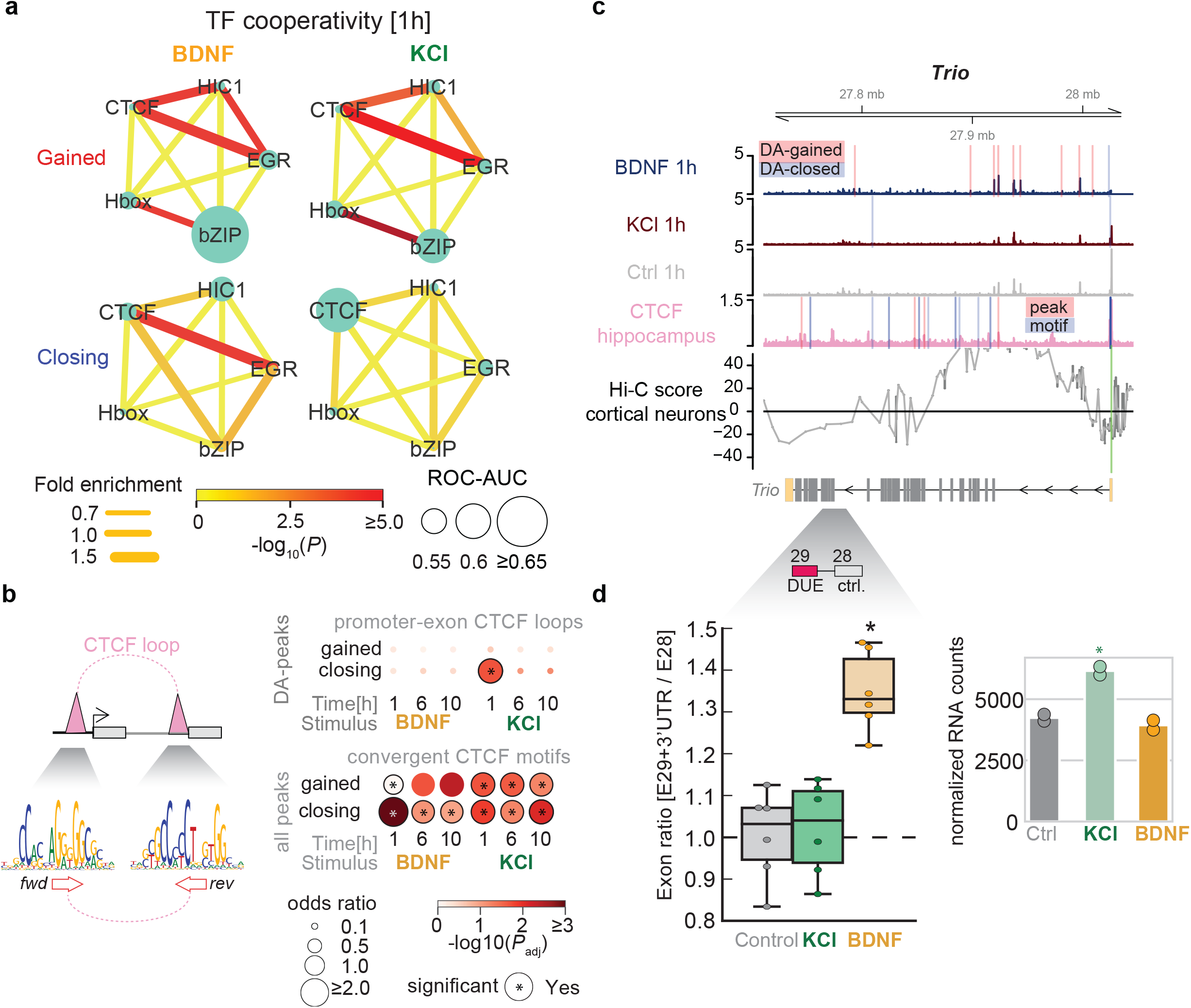
Chromatin-TF interactions during mouse neuronal activation and their association with promoter-exon loops and splicing. (a) Enrichment associations network between HIC1/bZIP/EGR/CTCF and Hbox based on results from (Fig 4a). Circle sizes indicate ROC-AUC using motifs for each TF alone in those peaks. Edges weights and colors indicate fold enrichment for co-occupied peaks versus single peaks, and significance of association. Calculations done with SuperExactTest(Wang *et al*, 2015). (b) (*top-left*) Scheme depicting CTCF-loop connecting promoters and exons. (*bottom-left*) CTCF-loops are enriched for convergent CTCF motifs. (*top-right*) Enrichment of promoter-exon CTCF loops in gained and closing DA-peaks (loop annotations from Ruiz-Velasco *et al*. (Ruiz-Velasco et al, 2017)). (*bottom-right*) Enrichment scores for convergent CTCF motifs in gained DA-peaks, closing DA-peaks and unchanged peaks associated all DA-peaks with a distance less than 50 Kbp. (c) Genome tracks harboring the *Trio* gene. ATAC-seq tracks indicate DA-peaks (red highlight = gained DA-peak; blue highlight = closing DA-peak); CTCF tracks indicate the presence of motifs (pink highlight = ChIP-seq peak; blue highlight = motif based on CTCF Position Weight Matrix). Below gene models, reference DUE exon position (red) is highlighted, and next it a block indicating a control exon (gray) used for comparison is highlighted. (d) (*left*) Exon ratio between E29+3’UTR and E28 fold changes 1h after treatment with BDNF (orange), KCl (green), and control (gray). Asterisk indicates significant changes versus control (*t*-test, two-sided; *P* < 0.1). (N = 2, independent biological replicates). (*right*) Normalized gene counts for gene expression values versus control (* = adjusted *P*-value < 0.1 versus control).

To test this idea, we quantified differentially used exons (DUEs) between BDNF and KCl, and found 54 genes with DUEs that harbored BDNF DA-peaks over a CTCF motif or a promoter-exon CTCF loop (OR=1.6 relative to non-DUE genes; *P* < 0.001; **Supplementary Figure 7b**). We selected DUEs within three of these 54 genes (Trio, Stxbp5, Cpe-201) whose functions were implicated in neurons (Katrancha *et al*, 2019; Woronowicz *et al*, 2010; Geerts *et al*, 2017) validated them using RT-qPCR. Relative exon usage was assessed by the ratio between the exon differentially used in our analysis and an one control exon from the same gene that remained unchanged after stimulation. We confirmed that BDNF but not KCl increased the relative exon usage of all three genes with respect to unstimulated control neurons without changing the expression level of the genes (**Fig. 5c-d** for Trio**; Supplementary 7c** for Stxbp5, Cpe-201; **Supplementary Data 3**). These results suggest that BDNF stimulation increased the expression level of specific spliced mRNA isoforms of some neuronal genes, whereas KCl stimulation did not exert this effect, probably through promoter-exon CTCF-looping regulation, which was described as splicing mechanisms before (Ruiz-Velasco *et al*, 2017). In the Trio gene DUE arises through inclusion/exclusion of exon 29, which is the last exon of a transcript variant that carries an additional 3’UTR sequence, generating a truncated protein. Given that reduced levels of full-length Trio and the truncated mutations are linked to neurodevelopmental disorders and intellectual disability (Pengelly *et al*, 2016; Sadybekov *et al*, 2017), our results highlight a potential connection between BDNF-induced chromatin dynamics and exon-specific gene expression in neuronal disorders.

### Chromatin accessible regions associated with neuropsychiatric traits

To investigate the relationship between activity-induced chromatin accessibility in neurons and human disease, we analyzed GO terms associated with the genes neighboring DA-peaks after BDNF and KCl stimulation and found they were linked with distinct neurobiological and learning functions (**Supplementary Figure 8a**). To determine whether these BDNF- and KCl-responsive elements are involved in distinct neuropathological traits, we used data from 45 genome-wide association studies (GWAS) that link genetic variants with complex traits of diseases (Buniello *et al*, 2019) and calculated the enrichment of trait-associated SNPs among our DA-peaks (transferred to the human genome) using partitioning heritability analysis (Finucane *et al*, 2015) (**Methods, Supplementary Data 4**).

We found enrichment for several neuronal traits for our full set of ATAC-seq peaks and little enrichment for non-neuronal traits (**Supplementary Figure 8b**), thus validating our analysis and experimental system. We identified 31 significant GWAS signals (adjusted *P* value < 0.1, using BH correction) in chromatin accessible regions from mouse neurons, suggesting that conserved neuronal chromatin regions (between the mouse and human genomes) are linked to a subset of psychiatric traits (**Fig. 6a**), consistent with previous studies showing that accessible genomic regions in mammalian brains are linked to human neuropsychiatric disorders (de la Torre-Ubieta *et al*, 2018; Hook & McCallion, 2020).

**Figure 6.**
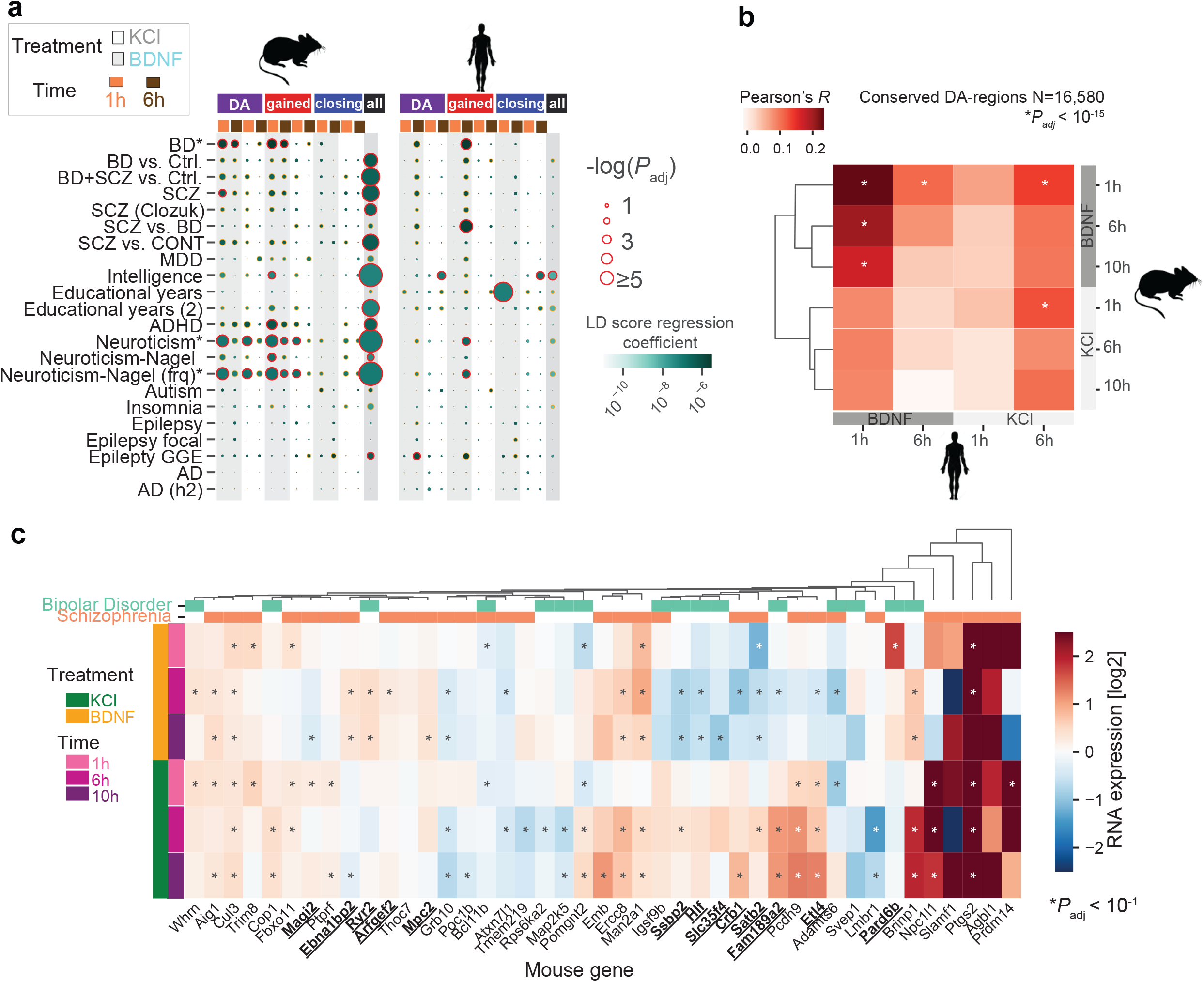
Conserved BDNF-specific chromatin accessibility associations with human complex traits. (a) Associations between chromatin accessibility DA-peaks and GWAS summary statistics. All DA-peaks (DA), gained DA-peaks (gained), closing DA-peaks, and consensus peaks (all) are fitted to summary statistics data of multiple GWAS studies (y-axis). Circle color indicates LDSC regression coefficient (effect size), and circle size indicates association significance. Shadings indicate treatments. Time points are shown (orange = 1h, brown = 6h). Red lines indicate LD score regression coefficient adjusted *P* < 0.1 after BH correction. Orange lines indicate *P* < 0.1. (BD = Bipolar Disorder; SZ = Schizophrenia; BD+SZ: BP and SZ samples combined; MDD = Major Depressive Disorder; ADHD = Attention deficit hyperactivity disorder; AD = Alzheimer’s Disease). Details for GWAS studies and preparation steps in Supplementary Data 4. (b) Correlations between differentially accessible elements conserved mouse and human regions across conditions (*N*=16,580). Asterisks indicate significance of Pearson’s *R* correlation after correction with BH-procedure. (c) Mouse expression of GWAS Schizophrenia and Bipolar Disorder human genes with DA-regions conserved and differentially accessible in the mouse genome. Genes with significant up- or down-regulation for one treatment (BDNF or KCl) and not the other are in bold and underlined.

In addition to these general enrichments, we also found several stimulation-specific trait associations. Specifically, gained DA-peaks after BDNF stimulation significantly overlapped with inherited risk loci for bipolar disorder (BD), Schizophrenia (SCZ), intelligence, attention deficit hyperactivity disorder (ADHD), and neuroticism (**Fig. 6a**). Among them, neuroticism risk loci were also associated with gained DA-peaks after KCl stimulation. DA-peak regions after BDNF and KCl did not overlap with risk loci for autism and epilepsy, implying that a subset of neuropsychiatric traits might be particularly sensitive to chromatin dynamics upon BDNF and KCl stimulation. Alzheimer’s disease (AD), a neurodegenerative trait, was not observed in our early stage neurons consistent with AD risk loci mainly being associated with enhancers in microglial cells (Hemonnot *et al*, 2019; Boix *et al*, 2021).

To validate the association between human neuropsychiatric traits and BDNF-induced DA-peaks in a different system, we generated human induced pluripotent stem cell (hiPSC) derived excitatory neurons, using doxycycline-inducible NGN2 expression, and determined chromatin accessibility after BDNF and KCl stimulation (**Methods**). The hiPSCs-induced neurons exhibited postmitotic neuronal markers comparable to the primary cultured neurons from mice (e.g., TUBB3, MAP2, PSD95 but not SYT1)(**Supplementary Figure 9a**). Similar to the results from mouse neurons, hiPSCs-induced neurons after BDNF stimulation generated more DA-peaks than KCl, although the major response was delayed compared to mice, being observed at 6h (**Supplementary Figure 9b-c**). Clustering of 3,029 human DA-peaks (FDR = 10%) revealed two separate groups by stimulation (**Supplementary Figure 9d**). A comparison of BDNF-induced chromatin accessibility between mouse and human neurons identified a weak but significant correlation in conserved genomic regions (**Fig. 6b**).

Repeating our partitioning heritability analysis using the human DA-peak regions (sorted by differential accessibility p-value) revealed 9 significant GWAS signals with neuropsychiatric traits (**Fig 6a; Supplementary Figure 8b**). A strong association with educational attainment was observed only in human DA-peaks upon BDNF stimulation. Conversely, the ADHD association found in mouse DA-peaks upon BDNF stimulation did not appear in human DA-peaks. However, BD, SCZ, and neuroticism were commonly associated with gained peaks after BDNF stimulation in both species but disappeared in overall accessible regions (**Fig. 6a**), demonstrating that neuropsychiatric disorder-related GWAS SNPs tend to be located near BDNF stimulation specific regulatory regions.

Prior work reported that individuals with BD or SCZ often show reduced levels of BDNF (Ray *et al*, 2014; Lima Giacobbo *et al*, 2019), suggesting a link between BDNF-mediated gene regulation and these traits. Therefore, we examined whether SCZ- and BD-risk loci-linked genes show an altered response upon BDNF stimulation that was different from their response upon KCl stimulation (which was not linked to BD or SCZ in our analysis in **Fig 6a**). Assessing the changes in expression levels of SCD- and BD-linked genes upon BDNF and KCl stimulation in mouse neurons (**Fig. 6c**) revealed BDNF-specific up-regulated (Ebna1bp2, Ryr2, Arfgef2, Pard6b, Mpc2) and down-regulated (Ssbp2, Hlf, Slc35f4, Crb1, Satb2, Fam189a2, Etl4, Magi2) DE-genes, implying that the GWAS-variants for these genes act through BDNF-response elements. Together, these analyses suggest a connection between activity-dependent chromatin accessibility and human complex traits, especially in a subset of neuropsychiatric disorders.

## Discussion

In this study, we performed a comprehensive analysis of gene expression and chromatin accessibility to define the overall principles and specificity of the response to neuronal activity triggered by BDNF, in comparison to neuronal activity triggered by KCl. Our analyses revealed that BDNF and KCl stimulation in mouse primary cortical neurons lead to distinct patterns of gene expression, involving early and late transcriptional waves, with differentially expressed TFs likely being responsible for the patterns of differentially expressed genes.

Regarding chromatin accessibility, we found that BDNF stimulation induced comprehensive changes in the enhancer landscape at an early stage, with concomitant gene expression changes, while KCl stimulation resulted in delayed chromatin remodeling of a similar set of enhancers. We speculate that the higher Fos protein levels in BDNF promotes more rapid opening of the distal regulatory regions, compared to KCl. Higher levels of Fos induction upon BDNF stimulation might be due to stronger activation of MAPK in BDNF compared to KCl, consistent with previous studies showing a correlation between MAPK activation and Fos expression (Whitmarsh, 2007). The variation in gene expression seen in BDNF and KCl likely arises because of the different levels and/or combinations of distinct co-factor TFs bound with Fos in their newly accessible regulatory regions. KCl-induced changes in chromatin accessibility in promoters at an early time-point showed little correlation with transcription, likely explained by more complex TF dynamics at promoters after KCl stimulation, involving TFs functioning as both activators and repressors (e.g., KLFs, E2Fs).

Fos, a classical pioneer factor assembling the AP-1 complex (Fos and Jun heterodimers, bZIP protein family members) (Biddie *et al*, 2011), was a major driver increasing chromatin accessibility for both stimuli. We further revealed that multiple co-regulatory TFs, such as ETSs, RFXs, and EGRs, are enriched with bZIP motifs and regulate gene expression, consistent with models that co-regulators recruited in the vicinity of bZIP TFs may provide functional specificity (Su *et al*, 2017). The association with EGRs and their higher expression in BDNF over time suggests a major involvement in the chromatin and expression events induced by BDNF, consistent with EGR TFs regulating downstream target genes involved in synaptic plasticity and memory formation (Beckmann & Wilce, 1997; Gallitano-Mendel *et al*, 2007). In addition, we found that the transcriptional repressor TF HIC1 (Pinte *et al*, 2004; Boulay *et al*, 2012; Ubaid Ullah *et al*, 2018) is associated both with subsets of binding regions opened by BDNF and with the overall closing of early accessible regions, correlating with its strong early up-regulation upon BDNF stimulation. The stronger co-enrichment of HIC1 and EGR in BDNF-gained peaks and weaker co-enrichment of CTCF and EGR in KCl-closing peaks (**Fig. 5a**) indicates a further regulatory role through interaction of these factors, as suggested previously (Pruunsild *et al*, 2017). Although we did not observe a clear role for HIC1 motifs in BDNF-induced Arc gene expression, HIC1 might act to regulate other genes as an early repressor in regions opened by bZIP and rapidly co-regulated by EGR factors. As co-binding of TFs may target a more specific set of genes and thus lead to a more specific functional impact (Jolma *et al*, 2015; Vandel *et al*, 2019; Ibarra *et al*, 2020), a complex interplay of these factors in the onset of BDNF specific gene expression is possible and requires further investigation.

The relationship between chromatin accessibility and genomic regions associated with complex traits can provide a deeper understanding for diseases in the context of genetic variation at non-coding regulatory regions (Maurano *et al*, 2012) that could affect TF occupancy (Vierstra *et al*, 2020). Previous studies showed that mapping mouse epigenomes with human conserved regions can reveal cis-regulatory regions linked to human GWAS (McClymont *et al*, 2018; Hook & McCallion, 2020). Here, we found that neuronal regulatory regions conserved between mouse and human are enriched for genetic variants linked to human neuronal traits, and more predominantly in regions that increase chromatin accessibility upon BDNF, suggesting that BDNF-activating chromatin associated with neuropsychiatric traits can be detected in several species. These associations with regions showing BDNF-induced gains of expression in both species highlight the utility of comparative genomics in prioritizing and validating associations with complex traits, with our study describing a systemic approach for dissecting stimulation-driven chromatin function in brain cells.

## Acknowledgments

We would like to thank the EMBL Sequencing Core Facility and the EMBL Proteomics Core Facility for help with sequencing and processing of samples related to this manuscript. We thank Daria Bunina, Mikael Marttinen, and Guy Riddihough for critical feedback of this manuscript, and Na Cai for input in our partitioning heritability analyses. I-YH was supported by a fellowship from the EMBL Interdisciplinary Postdoc (EIPOD) programme under Marie Sklodowska-Curie Actions COFUND (grant agreement number 664726). This work is supported by the DFG fund #SPP 1738 to K.M.N. and NIH grant # R01 HG010501 to M.L.B.

## Data availability

RNA-seq and ATAC-seq raw data for mouse and human, including normalized counts and processed genes and peaks comparison have been deposited in Gene Expression Omnibus, with accession code GSE169708. The mass spectrometry proteomics data have been deposited to the ProteomeXchange Consortium via the PRIDE (https://pubmed.ncbi.nlm.nih.gov/30395289/) partner repository with the dataset identifier PXD022378.

## Code availability

Processed data and scripts of relevant computational analyses are available at https://git.embl.de/grp-zaugg/neuronal_activity_bdnf.

## Author contributions

V.S.R and K.-M.N. conceived the study. J.B.Z and K.-M.N. jointly supervised the computational parts of the study. V.S.R. performed mouse primary neurons sample preparations, ATAC-seq, RNA-seq, CRISPR, and exon splicing studies under the supervision of K.-M.N. I.L.I. conducted all pre- and post-processing computational analyses, under the supervision of J.B.Z. L.G. conducted proteomic sample preparation and data interpretation from mESC-derived neurons, and RT-qPCR measurement and interpretation for enhancer CRISPR knockouts, under the supervision of K.-M.N., L.M., K.W. and M.L.B. prepared a database of mouse chromatin negative regions for motif enrichment and helped in the interpretation of 8-mer enrichment results. I.H and M.S. helped with data acquisition. H.M.H conducted proteomics measurements and post-processing analyses under the supervision of M.M.S. I.L.I. and V.S. wrote the original manuscript with input from K.-M.N., and J.B.Z. I.L.I, K-M.N. and J.B.Z wrote the manuscript, with the input and approval of all co-authors.

## Supplementary Information

**Supplementary Figure 1.**
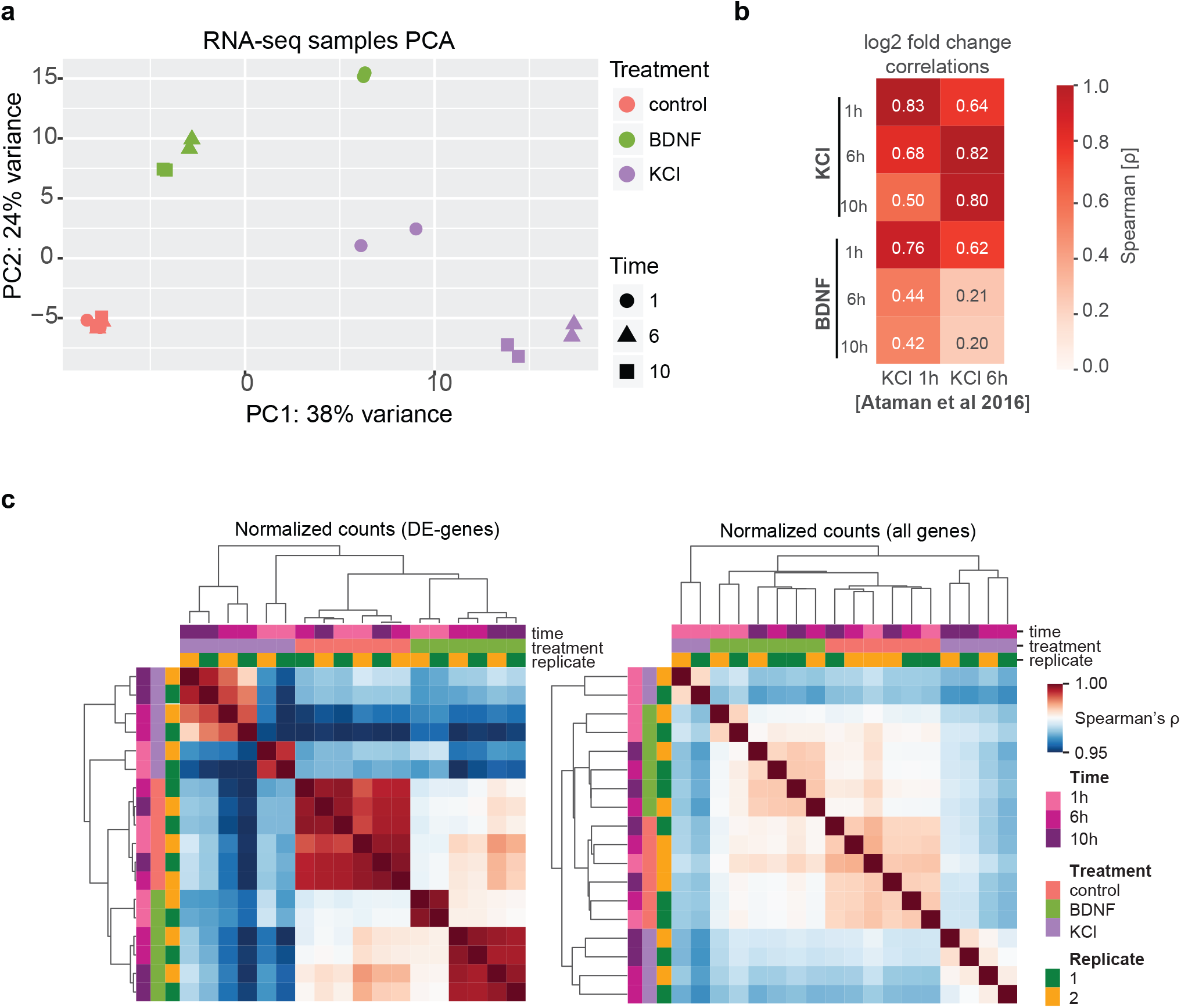
RNA-seq computational data processing. (a) Principal component analysis visualization RNA-seq samples based on DESeq2 normalized counts (treatment = colors; time=shapes). (b) Gene expression Log2 fold change correlations between genes from BDNF- and KCl-stimulated neurons (y-axis), versus DE-genes reported in mouse cultured neurons stimulated with KCl (x-axis) (Ataman *et al*, 2016). (c) (*left*) Hierarchical clustering of all RNA-seq samples, using the union of DE-genes found in any comparison versus control. (*right*) Hierarchical clustering of all RNA-seq samples, using all genes and normalized counts.

**Supplementary Figure 2.**
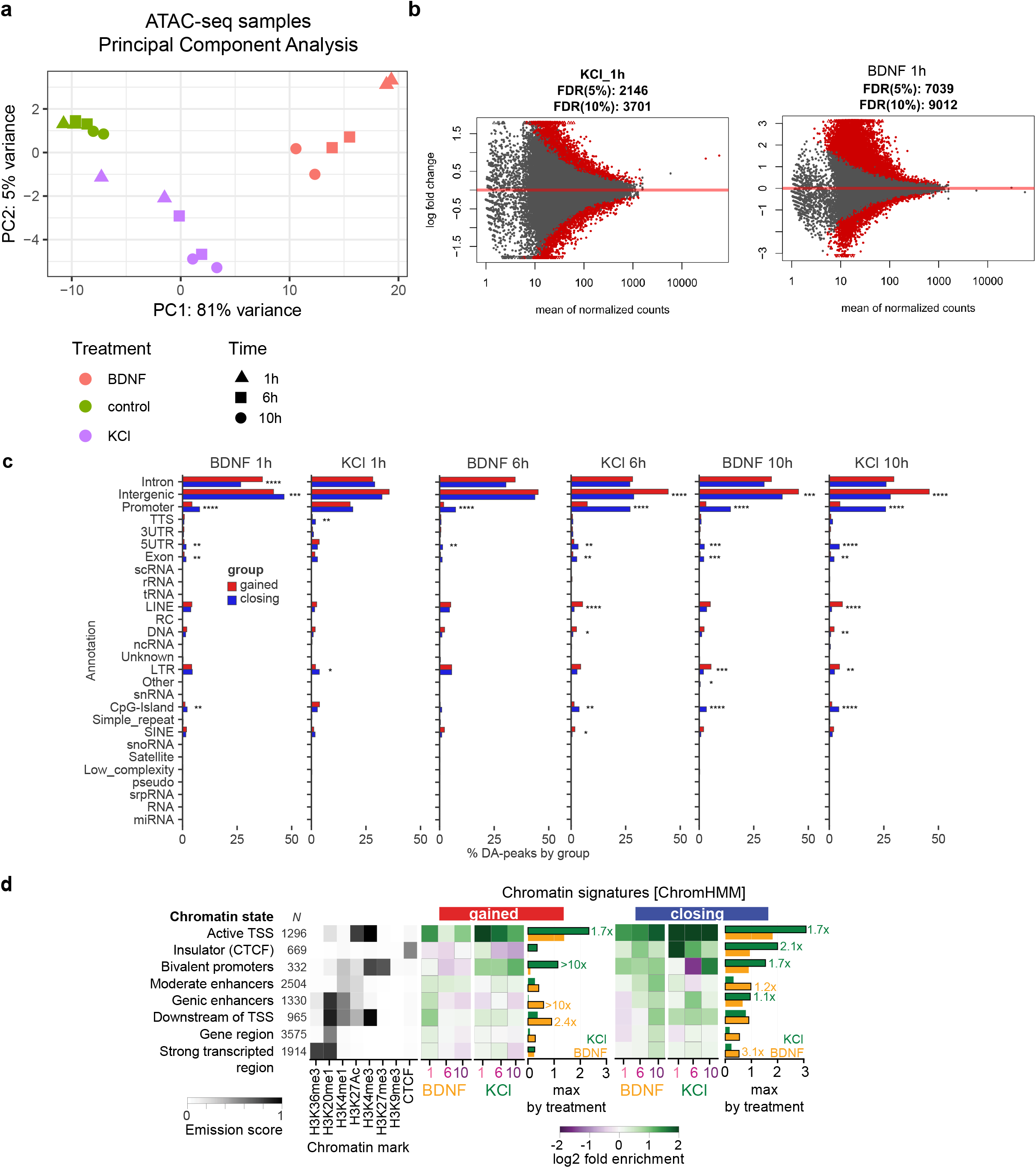
ATAC-seq computational data processing. (a) Principal component analysis visualization for treatment and time points using normalized counts generated with DiffBind [**Stark and Brown, 2011. Bioconductor**]. (b) MA-plots for consensus peaks. Differentially accessible peaks (DA-peaks; red dots) obtained in ATAC-seq samples for KCl 1h (left) and BDNF 1h (right). Red points indicate a peak with accessibility levels higher (log fold change > 0) or lower (log fold change < 0) compared to matched controls (FDR=10%). (c) Percentage distribution of DA-peaks for BDNF and KCl 1h in peak annotations (using HOMER). Asterisks indicate within-stimulation Fisher’s exact test comparisons, if annotation fraction are statistically significantly different between gained and closing DA-peaks (*adjusted *P* <0.05; \*\**P* < 0.01; \*\*\**P* < 0.001). (d) Chromatin states enrichments in DA-peaks for neuronal marks, based on ChromHMM 15-states model using neuronal epigenomics datasets (**Methods**). *N* indicates the total number of DA-peaks associated with each annotation. Black heat map indicates emission probabilities. Green/purple heat maps indicate fold enrichment between nucleotides coverage for DA-peaks in those annotations and background. Black-contoured squares indicate change with the highest log2 fold change in each stimulation. Barplots summarize the highest log2 fold enrichment value for BDNF (yellow) and KCl (green). Fold enrichments with greatest change between BDNF and KCl highest values are labeled.

**Supplementary Figure 3.**
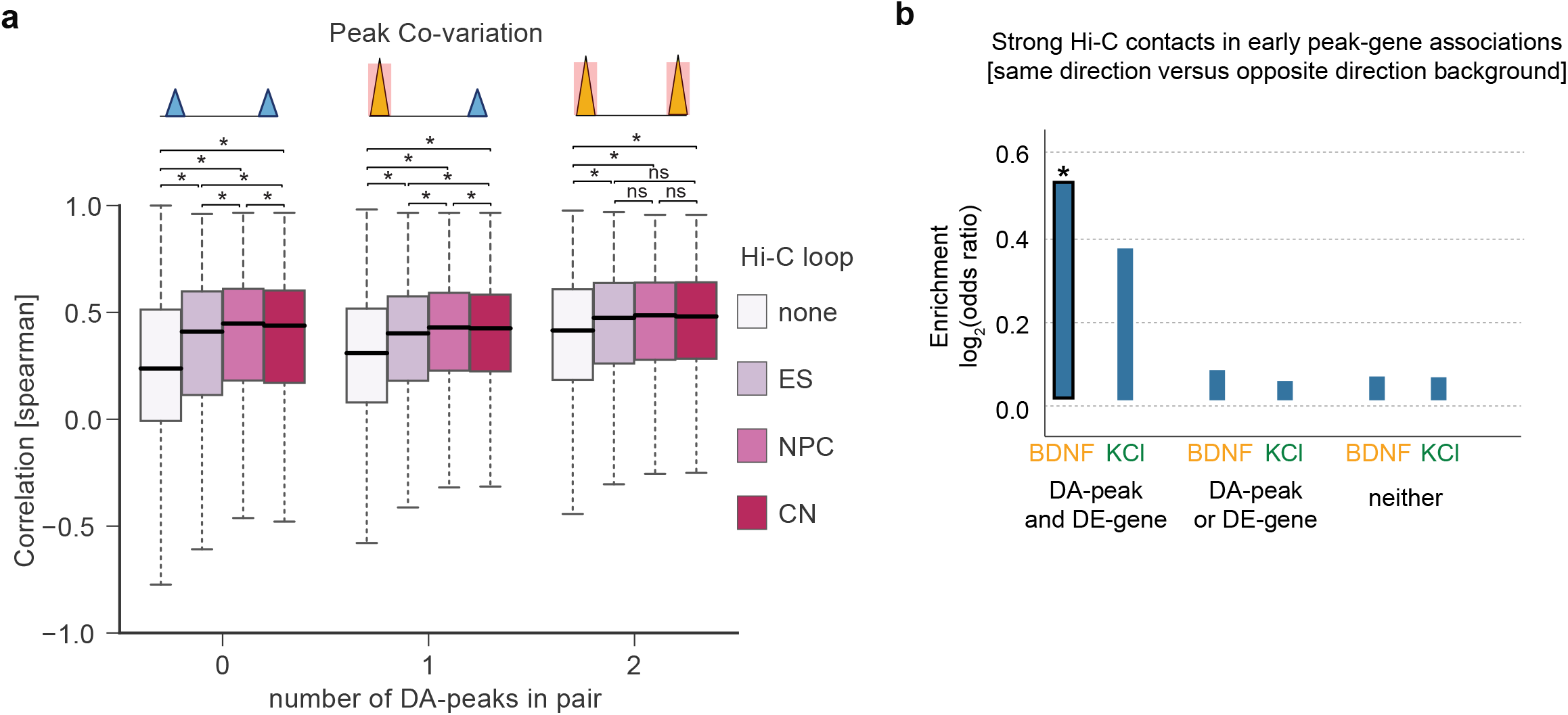
Chromatin peaks variability and association with High-conformation Capture data (Hi-C) (a) Correlation distributions for normalized counts of proximal peak pairs with zero, one or two DA-peaks in that pair, grouped by Hi-C data evidence (CN=cortical neurons; NPC=neural progenitor cells; ES=Embryonic stem cells). Hi-C data from (Bonev *et al*, 2017). (b) Enrichment of same sign log2 fold changes for pairs of DA-peaks and proximal DE-genes when grouped as positive Hi-C loops (strong Hi-C contacts), versus different signs in the same group. Asterisk indicates P < 0.1 significant enrichment based on Fisher’s exact test (BH adjusted).

**Supplementary Figure 4.**
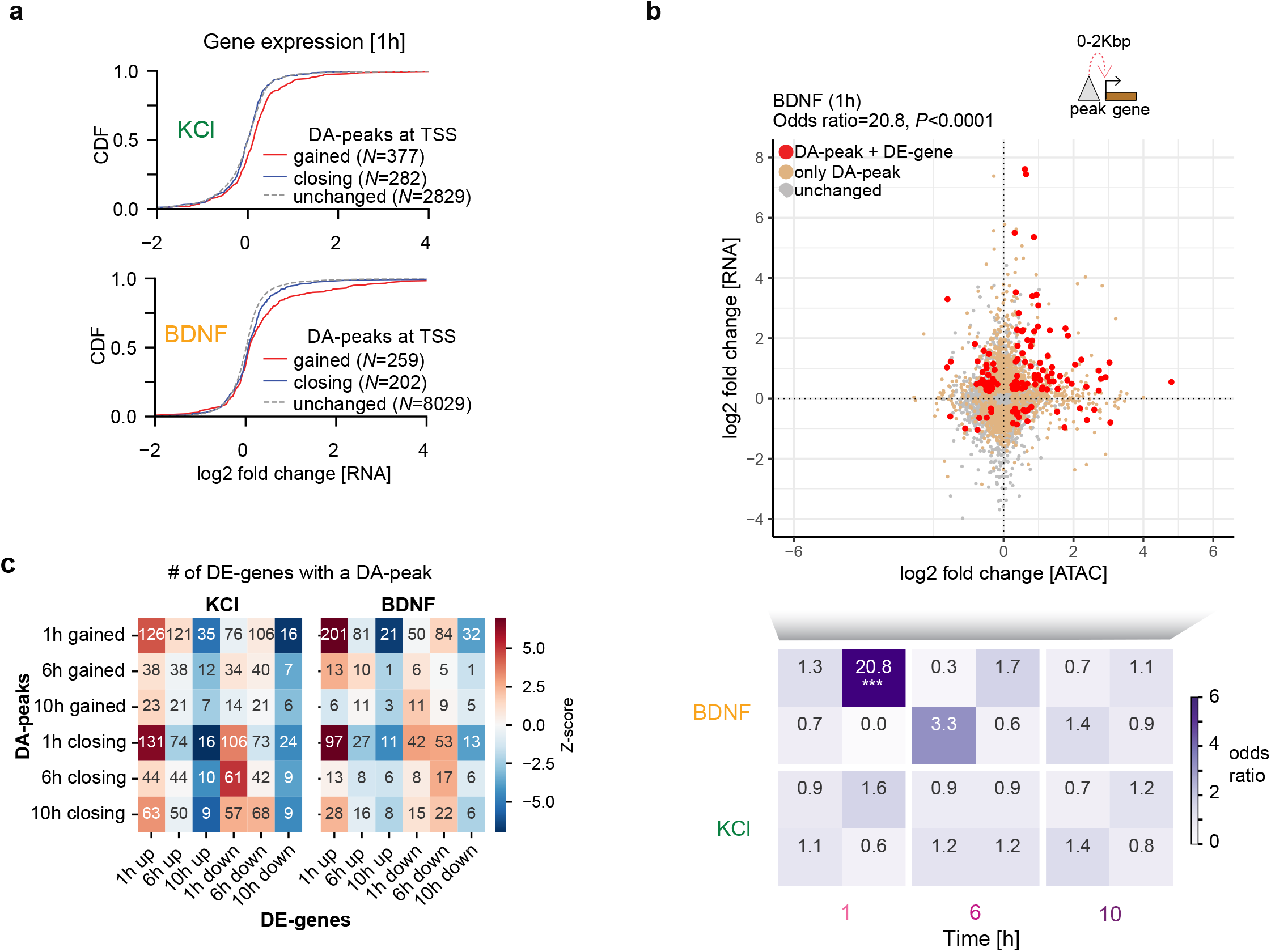
Association between DA-peaks and genomic and RNA features. (a) Cumulative distribution function of expression changes (log2) between genes based on the presence of DA-peaks (gained or closing), or unchanged (gray line). (b) (*top*) Association between peaks at transcription start sites (TSSs) and gene expression at BDNF 1h. Each point indicates the log2 fold change of an ATAC-seq peak and the gene expression of a closest gene with distance between 0-2Kbp from TSS regions. Colors indicate whether none (gray), only the peak (orange), or both peak and gene (red) show significant changes versus control. (*bottom*) Enrichment for paired DA-peak and DE-gene in the four quadrants are summarized for BDNF and KCl. Asterisks indicate *P* values as corrected by a Benjamini Hochberg procedure (* = *P* < 0.05; ** = *P* < 0.01). (c) Number of associations found between DE-genes comparisons (x-axis) and DA-peaks comparisons (y-axis) based on same (up/gained; down/closing) or opposite directions. Colors indicate significance Z-scores based on observed versus expected values from permutations.

**Supplementary Figure 5.**
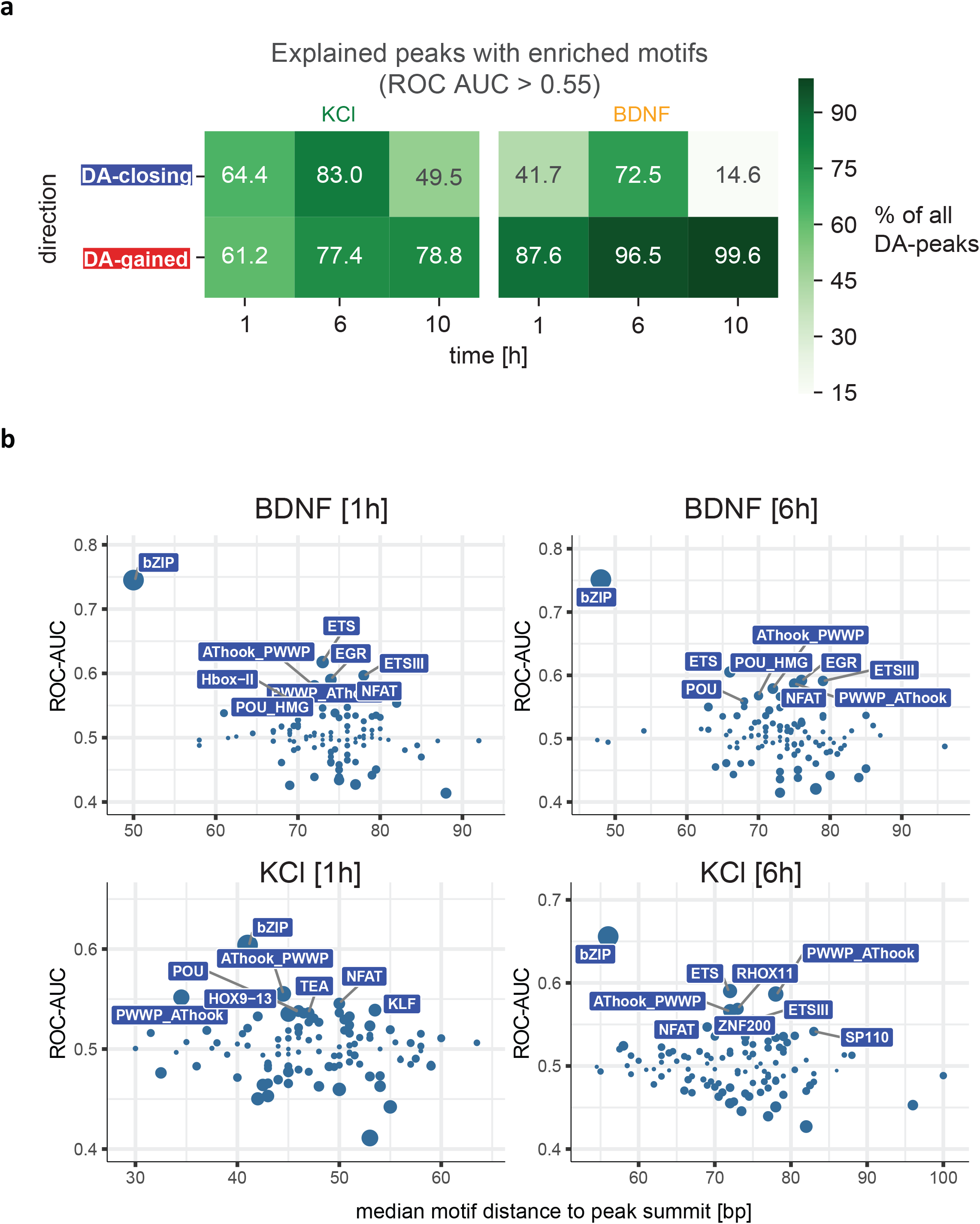
Enrichment of TF motifs in BDNF and KCl differentially accessible ATAC-seq peaks. (a) Total fraction (as percentage) of DA-peaks explained by enriched TF motifs (minimum ROC-AUC > 0.55). (b) *8*-mers modules harboring gained DA-peaks are visualized by its overall enrichment (area under the receiver operating characteristic curve, ROC-AUC, *y*-axis) versus median relative distance to the peak summit for all motif observations (*x*-axis). bZIP has the highest enrichment and additionally the lowest median distance to summit for BDNF 1 and 6h.

**Supplementary Figure 6.**
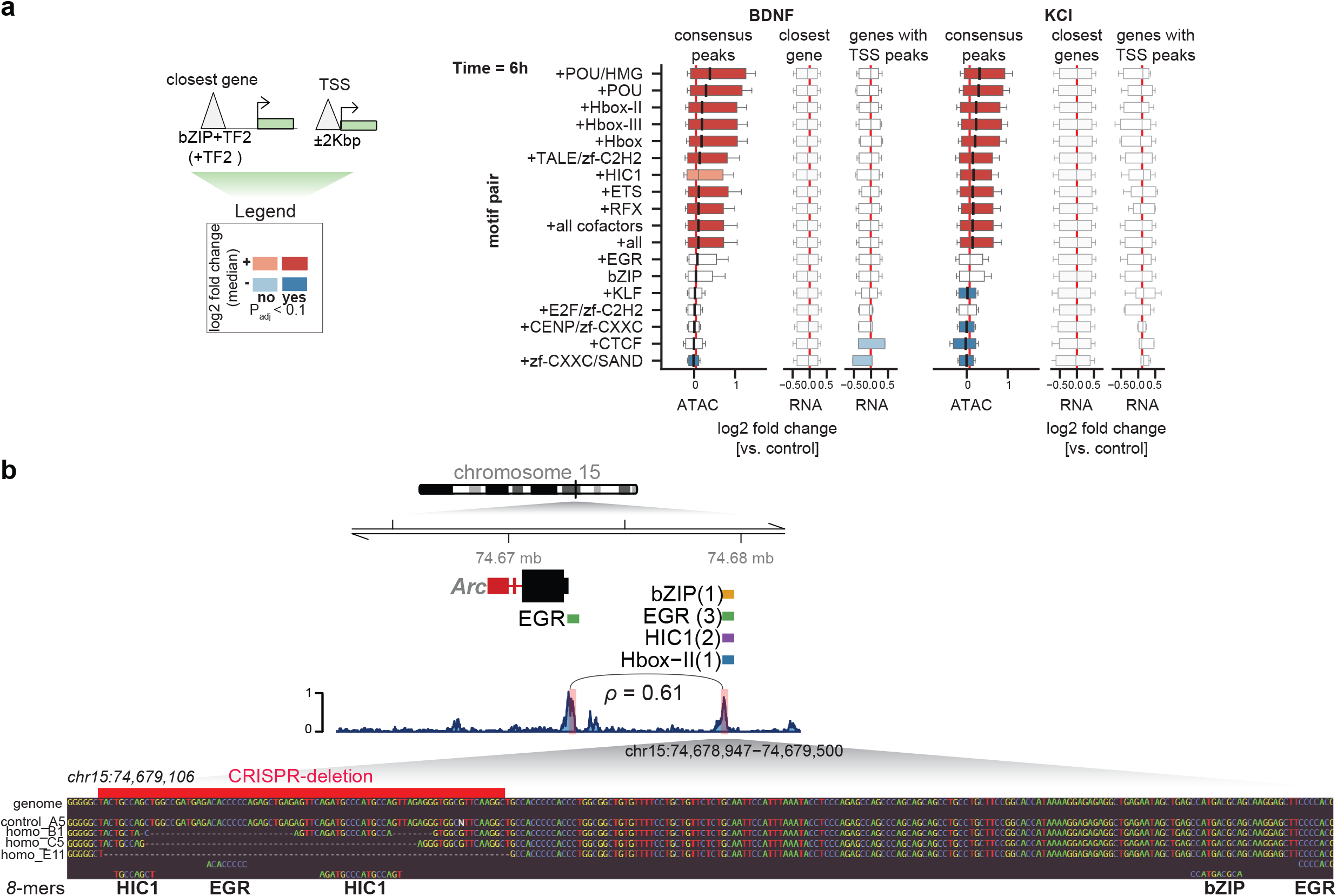
Effect of bZIP and co-regulatory TFs on BDNF and KCl at 6 hours post stimulation, and genome tracks for modified Arc distal regulatory element perturbed using CRISPR in mESC-derived neurons. (a) Changes in log2 fold change between bZIP containing peaks and bZIP+TF2 containing ATAC-seq peaks in time point 6h. Legends as in Figure 4a-b (b) (*top*) Genome track for Arc gene and distal enhancer. Highlighted mutated enhancer region in gray box. Coordinate indicates consensus peak region using DiffBind. Numbers in parentheses indicate the absolute number of motifs detected for each highlighted TF specificity module. (*bottom*) aligned nucleotide sequences for one control replicate (A5) and three homozygous replicates (B1, C5, E11). aligned in the mouse genome (mm10). Red region indicates the deleted region based on sequencing data. Motifs for ERG, bZIP, HIC1 and Hbox are highlighted whenever a region with *8*-mer hits with a high score is detected.

**Supplementary Figure 7.**
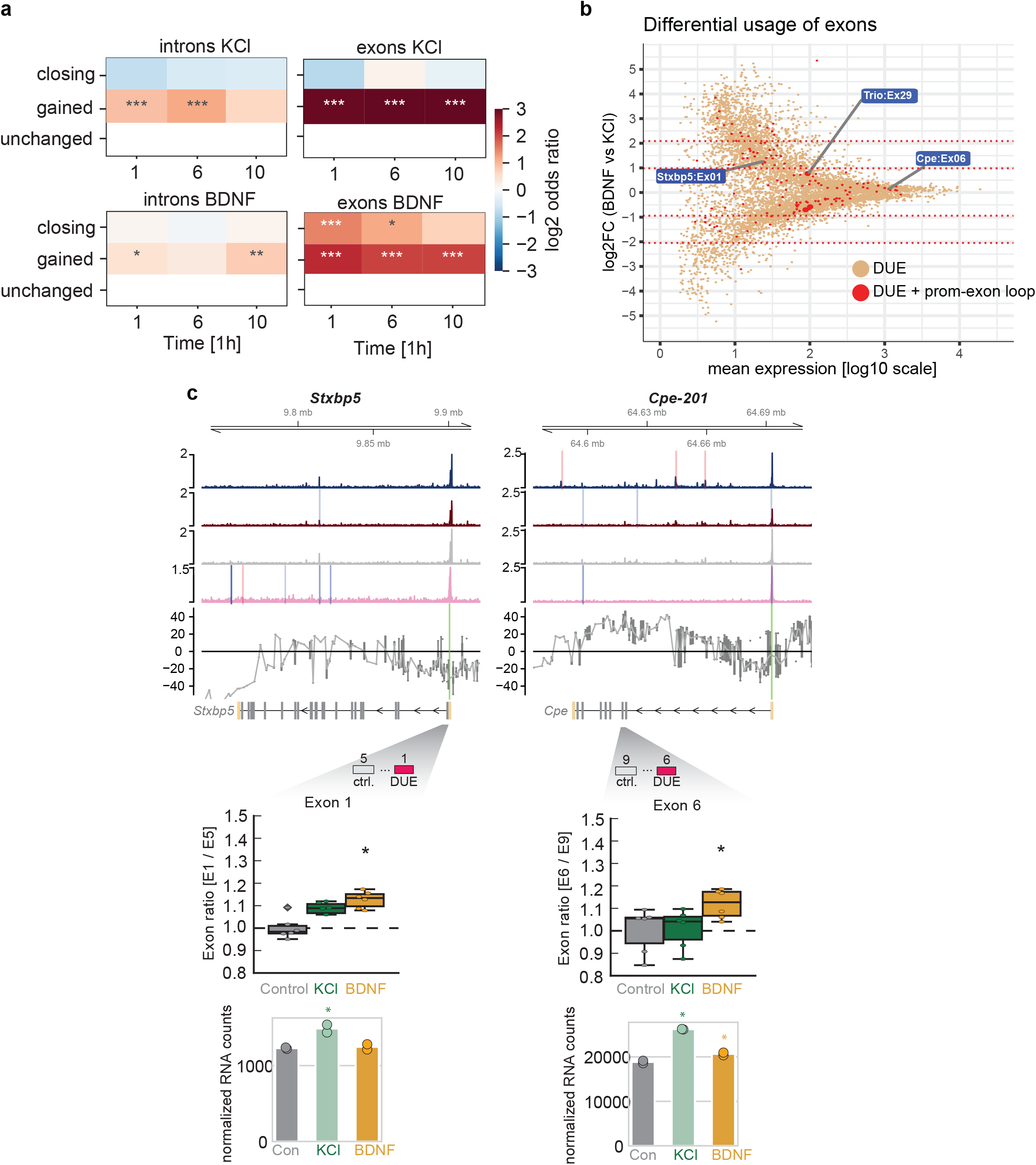
Association between promoter-exon CTCF-loops, chromatin DA-peaks and differentially usage of exons. (a) Enrichment scores for convergent CTCF motifs in gained DA-peaks, closing DA-peaks and unchanged peaks associated with introns (*left*) and exons (*right*). Annotations are based on HOMER. Asterisks indicate Fisher’s exact test *P* value significance, adjusted by Benjamini Hochberg procedure (*, **, *** = P < 0.05, 0.01 and 0.001, respectively). Circle color and size annotations as in (b) (b) Exon log2 fold changes between BDNF and KCl as quantified by DEXSeq(Anders *et al*, 2012) (**Methods**). Dots indicate differentially used exons (orange), differentially used exons with promoter-exon CTCF loops (reds) or unchanged exons (gray). Labels indicate genes and exon names with both features, selected for experimental validation (c) *(top)* Genome tracks for *Stxbp5* and *Cpe201*. ATAC-seq tracks indicate DA-peaks (red = gained; blue = closing) and CTCF tracks indicate presence of motifs (pink = ChIP-seq peak; blue = motif). Below gene models, reference DUE exon position (red) is highlighted, and next it a block indicating a control exon (gray) used for comparison is highlighted. *(middle)* Box plots indicate fold change ratios between reference exon and control exon fold changes 1h after treatment with BDNF (orange), KCl (green), and control (gray). Asterisk indicates significant changes versus control (adjusted *P* < 0.1; *t*-test, two-sided). (*bottom*) Normalized RNA counts (asterisk = significant log2 fold change versus control).

**Supplementary Figure 8.**
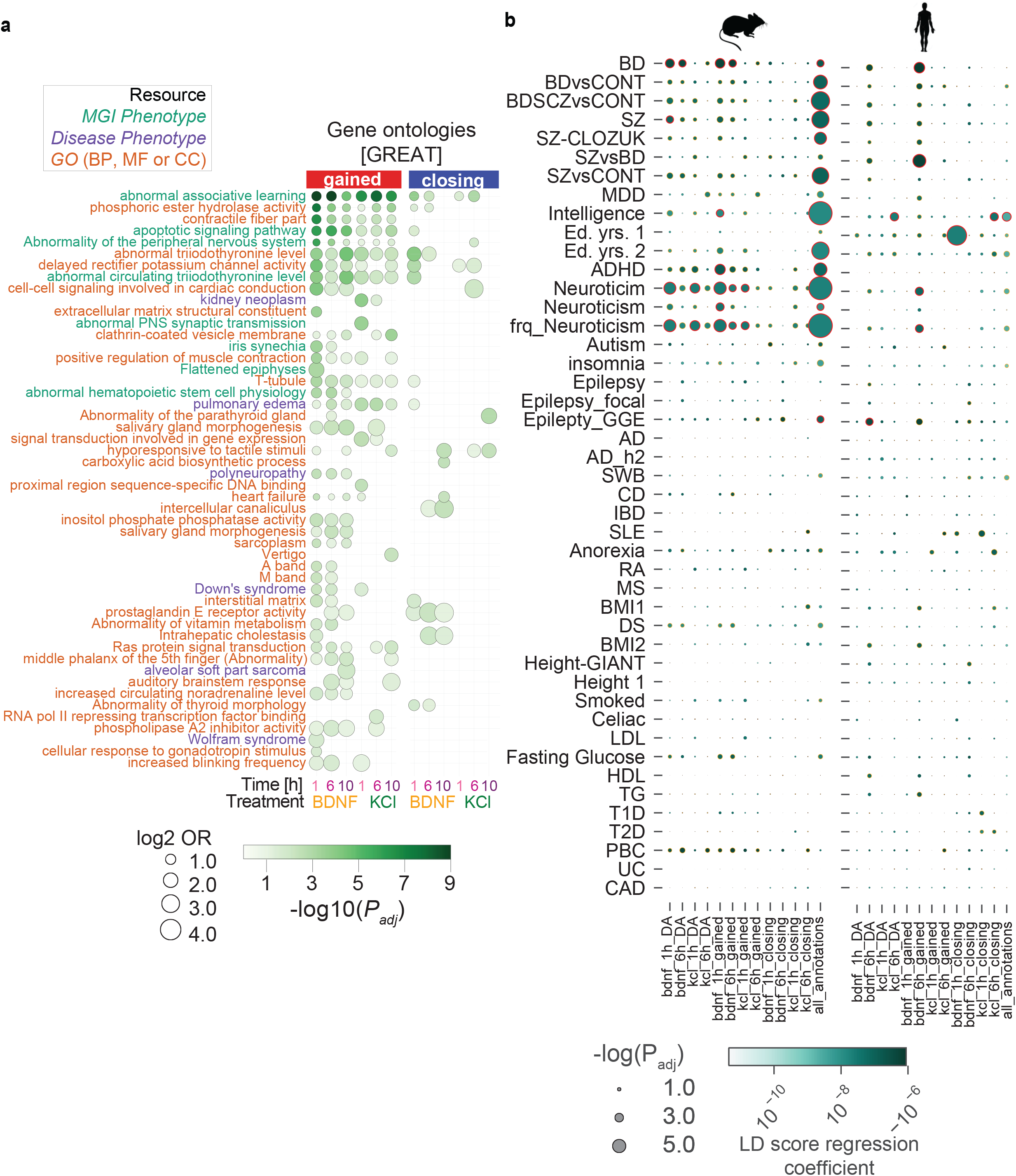
Disease Ontology associations for mouse DA-peaks and Partitioning heritability analysis of mouse primary cortical neurons and hiPSC-derived neurons. (a) Genes Ontology enrichments for genes associated with DA-peaks, linked based on GREAT (McLean *et al*, 2010). Circle size indicates effect size (log2 Odds Ratio). Color indicates significance. *Y*-axis color labels indicate source of ontology. (b) Associations between mouse consensus and DA-peaks from cortical neurons and human GWAS terms based on summary statistics. Legends as Figure 5a. GWAS traits labeled as associated with neuronal function are on top of the heatmap. GWAS related to non-neuronal traits are at the bottom of the heatmap.

**Supplementary Figure 9.**
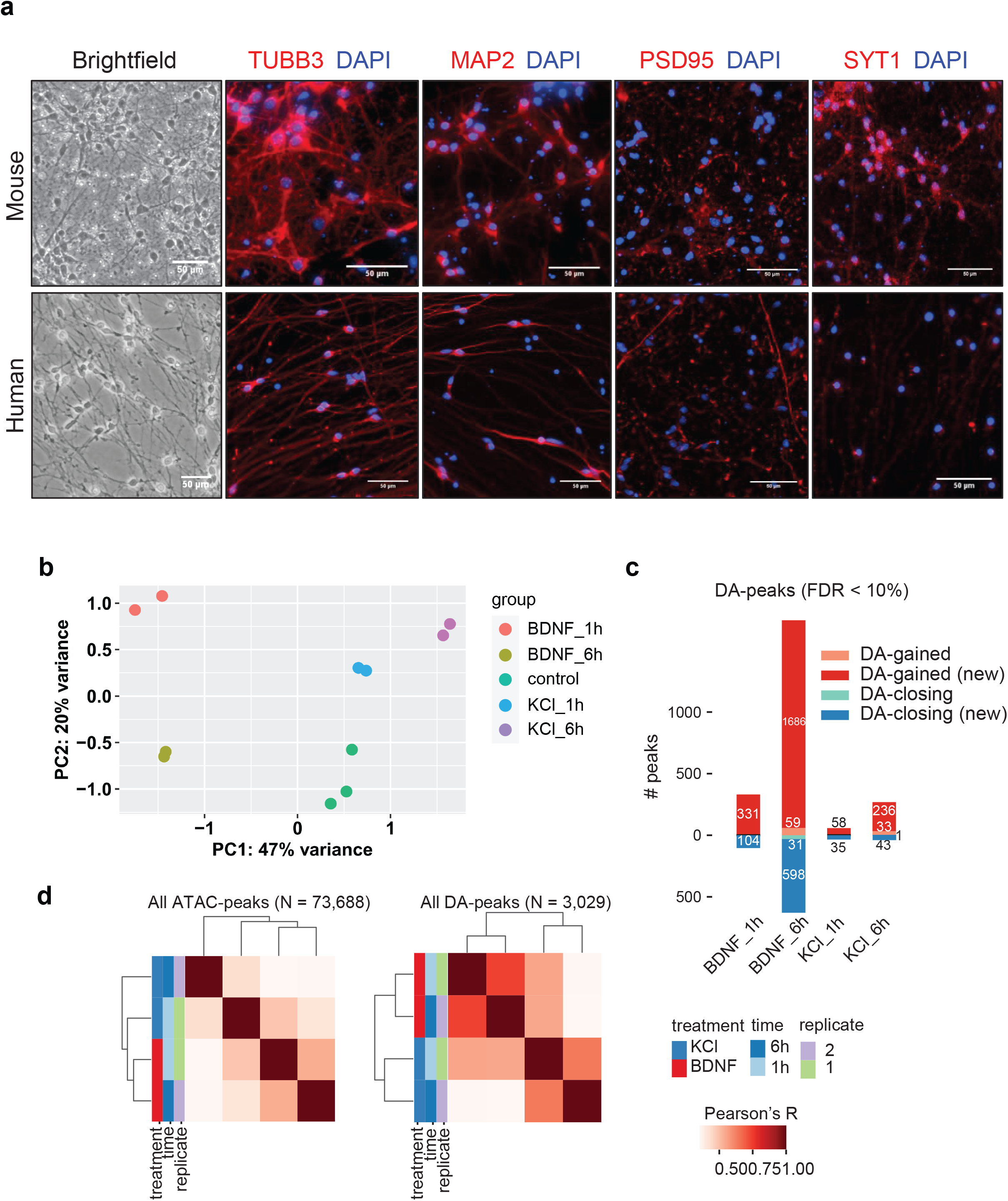
Histological staining of mouse and hIPSC-induced neurons and differentially accessible peaks upon treatment with BDNF and KCl. (a) neuronal maturation markers of cultured mouse cortical neurons (DIV7) (*top*) and hIPSC-derived neurons (*bottom*). Bright-field and merged fluorescence images of neuronal markers, β-tubulin III (TUBB3) and Map2 (MAP2), and synaptic proteins, postsynaptic density protein 95 (PSD95) and synaptotagmin I (SYT1), with DAPI (nuclei). Scale bar = 50 µm. Visualized dendritic branches and the presence of synaptic markers are characteristic of functional mature neurons. Scale bar = 50 µm. (b) Principal component analysis visualization of chromatin accessibility changes in hIPSC-induced neurons. (c) Number of DA-peaks recovered for each time point and stimulation category when compared versus control hIPSC-derived neurons. Gained (new) and closing (new) indicate peaks observed as differentially accessible for the first time in that time point, whereas gained and closing indicate DA-peaks already observed as DA in a previous time point. (d) Hierarchical clustering of normalized counts obtained from ATAC-seq consensus peaks (*left*) and only DA-peaks (*right*).

## Methods

### Primary cortical neuron culture

Prenatal embryos of CD-1 mouse at embryonic day 15 (E15) were used for the isolation of cortical neurons. Embryonic cortex was isolated and dissociated by chopping with scalpel followed by digestion in Accutase (ThermoFisher, A1110501) for 12 mins at 37°C. During digestion we treated the tissue with 250 unit/µL of Benzonase (Millipore, 71206-3) to prevent neuronal clumping due to genomic DNA released from dead cells. Following digestion, neurons were triturated gently and passed through the 40µm cell strainer (BD Falcon, 352340) before plating them onto a 6-well plate at a density of 1×10^6^ cells per well. Tissue culture plates were coated with 0.1 mg/mL of Poly-D-Lysine (Sigma, P0899) and 2.5 μg/mL of laminin (Sigma, 11243217001). Primary neuronal cultures were maintained in Neurobasal medium (ThermoFisher, 21103049) containing 1% penicillin/streptomycin (ThermoFisher, 15140122), 1% GlutaMAX (ThermoFisher, 35050), and 2% B27 supplement (ThermoFisher, 12587010) at 37°C with 5% carbon dioxide in the incubator. Post-seeding after 1 day in vitro (DIV1), half of the media was replaced with fresh pre-warmed Neurobasal media with all the supplements.

### Neuronal differentiation of mESCs

Differentiation of mouse embryonic stem cells (mESC) to glutamatergic neurons was performed as previously described (Bibel *et al*, 2007) with minor modifications. Briefly, we removed the leukemia inhibitory factor (LIF) from mESC culture and grew cells in suspension using non-adherent plates to enhance the formation of embryoid bodies. Every two days, differentiation medium without LIF was exchanged and on day 4, 5 μM of retinoic acid (Sigma, R26250) was added to the medium to promote the differentiation to the neuronal lineage. On day 8, neuronal precursor cells (NPCs) were dissociated using trypsin and plated at a density of 2×10^5^ cells per cm^2^ on plated coated with 0.1 mg/mL of Poly-D-Lysine (Sigma, P0899) and 2.5 μg/mL of laminin (Sigma, 11243217001). Cells were cultured on N-2 medium, consisting on regular DMEM supplemented with 1x N-2 supplement (Thermo Fisher Scientific #17502048), 1x B27 supplement (Thermo Fisher Scientific #12587010) and antibiotics), which was replaced every two days. Experiments were always performed 4 days after plating cells.

### Neuronal differentiation of hiPSCs

Human iPSCs derived from peripheral blood mononuclear cells, kindly provided by Michael Snyder lab in Stanford University, were cultured in Essential 8 medium with supplement (ThermoFisher, A1517001) and vitronectin (ThermoFisher, A14700) coated dish. iPSCs were transduced with hNGN-2 and rtTA lentivirus at 60% of confluency. After 2 days of transduction hNGN2-eGFP media was replaced with 1:1 ratio of Neurobasal A (ThermoFisher, 10888022) and DMEM/F12 medium containing GlutaMAX, Insulin (Gibco, 51300-044), N2/B27 without Vitamin A (ThermoFisher, 17502048/12587010) supplements with 30% of glucose together with 1μg/ml of doxycycline to induce rtTA expressions (Day 0). Then GFP-positive hiPSCs were selected by adding 3 µg/ml puromycin (Day 2). After 2 days of puromycin selection, cells were dissociated with Accutase (ThermoFisher, A1110501) and replated in triple coated dish with 0.1 mg/mL of Poly-D-Lysine (Sigma, P0899), 10μl/ml of Laminin (Sigma, 11243217001), and 10μl/ml of Fibronectin (Sigma, F0895) in Neurobasal medium (ThermoFisher, 21103049) containing B-27 w/o Vitamin A (Gibco, 12587010), GlutaMAX, 200µM of L-Ascorbic Acid (Sigma, A5960) and 1µM of DAPT (Tocris, 2634) (Day4). 1µg/μl of puromycin was added for 2 more days and 1µg/μl doxycycline for 4 more days. Media was replaced every 2-3 days. After 10 days (Day14), iNeurons are collected and subjected to ATAC-seq.

### Lenti-virus generation

Lenti-viruses each containing hNeurogenin-2 (hNGN-2) under the control of TetON promoter and reverse tetracycline activator (rtTA) was used (Zhang et al., 2013 PMC3751803). Lenti-virus was generated by co-transfecting with two helper plasmids, pMD2 and psPAX2, by calcium phosphate transfection with CalPhos™ Mammalian Transfection Kit (Takara, 631312) following the manufacturer’s protocol into HEK293T cells. After 48-72hr of incubation media containing lentiviruses were harvested and concentrated at 28.000 g for 3 hours, aliquoted and stored at - 80°C.

### Immunofluorescent staining of mouse neurons

Mouse primary cortical neurons on DIV7 were fixed on coverslips with 3% para-formaldehyde solution in PBS for 20 minutes at room temperature. Fixation was quenched with 30 mM glycine for 5 minutes and cells were subsequently washed thrice in PBS. Cells were permeabilized with 0.1% Triton-X 100 for 5 minutes, washed again thrice with PBS and blocked in 0.5% BSA for 30 minutes at room temperature. Primary antibodies (mouse monoclonal anti-Map2 (Sigma, m9942), rabbit polyclonal anti-Synaptotagmin (Cell Signaling Technologies, 3347), mouse monoclonal anti-beta III Tubulin (Abcam, ab78078) and mouse monoclonal anti-PSD95 (Thermofisher, MA1-046)) were diluted 1:200 in 0.5% BSA (with the exception of Synaptotagmin, 1:50) and incubated for 1h at room temperature, followed by washes with PBS before the addition of the secondary antibody (goat anti-rabbit or mouse IgG conjugated to Alexa Fluor 594, (Life Technologies, A11012 and A11005), diluted 1:500 in 0.5% BSA, for 30 minutes at room temperature. Nuclei were stained with 5 µg/ml DAPI and samples were washed thrice with PBS. Coverslips were mounted on slides with Prolong Gold antifade reagent (Life Technologies). Images were taken with a Nikon Ti-E widefield microscope and processing of images was done in Fiji.

### Stimulation with BDNF and KCl

Prior to every stimulation on DIV7, neurons were made quiescent for 2 hours with 100 μM D-2- amino-5-phosphonopentanoic acid (D-AP5; Tocris #0106) and 1 μM tetrodotoxin (TTX; Tocris #1078/1). KCl (55mM) depolarization was performed by adding warmed KCl depolarization buffer (170 mM KCl, 2 mM CaCl_2_, 1 mM MgCl2and 10 mM 4-(2-hydroxyethyl)-1- piperazineethanesulfonic acid (HEPES)) to a final concentration of 31% directly into the neuronal culture medium and incubated for 1, 6, and 10 hour. For the BDNF (R&D systems, 248-BDB) stimulation, neurons were incubated with BDNF (10 ng/ml) on DIV7 for 1, 6, and 10 hour.

### Deletion of *Arc* putative enhancer with CRISPR technology and determination of Arc expression by qPCR

To disrupt a putative EGR binding regions of the Arc *gene* enhancer, two guide RNA sequences (gRNA 5’ TTGGGGGGGCTACTGCCAGC 3’ and 5’ CTTGAACGCCACCCTCTAAC 3’) were cloned into pspCas9(BB)-2A-GFP (Addgene; #48138) and pspCas9(BB)-2A-RFP (Addgene; # 91854), respectively, following the protocol by Ran et al (Ran *et al*, 2013). Of each resulting plasmid, 2 μg were nucleofected into 2× 10^6^ mESC using a Nucleofector (Lonza). After 48 h, samples were single-cell sorted for GFP and RFP double positive cells. Colonies were expanded for genotyping and freezing. Deletion events were confirmed for homozygosity by agarose gel electrophoresis and checked by Sanger sequencing.

A total of three mutant homozygous lines and three CRISPR control lines -resulting from CRISPR/Cas editing rounds that did not include the desired deletion- were differentiated in duplicates to glutamatergic neurons and stimulated for 1h with BDNF or KCl. *Arc* expression upon stimulation was quantified by RT-qPCR. cDNA was generated from total RNA treated with DNase (Thermo Fisher) using MultiScribe^TM^ Reverse Transcriptase (Thermo Fisher). A total of 10 ng of cDNA were subjected to qPCR using PowerUp SYBR Green Master Mix (Applied Biosystems) and the StepOnePlus Real-Time PCR system. Each reaction was assayed in triplicates (BDNF stimulation) or duplicates (KCl stimulation). Changes in *Arc* expression were assessed by normalization to *Rpl-13* and unstimulated control (2^-ΔΔCt^).

### RNA-seq sample preparation

Mouse cortical neurons were collected at 1, 6, and 10 hours after each stimulation for RNA isolation. The RNeasy kit (Qiagen) was used to extract RNA and genomic DNA was digested using the Turbo DNAse kit (Ambion). To assess quality of RNA all samples were analysed using Bioanalyzer (Agilent Genomics). Only samples with a RIN (RNA integrity number) score above 9 were used for library preparation. To prepare libraries we used the oligo-dT capture kit (NEB) in combination with the NEBNext Ultra II kit. We pooled 24 samples with each sample carrying a distinct barcode and sequenced on NextSeq 500 at EMBL, Heidelberg Gene Core facility.

### RNA-seq computational data processing

We mapped RNA-seq reads in each sample to the *M. musculus* mm10 genome using TopHat2 (v2.1.1) (Kim *et al*, 2013), using the transcriptome defined in Gencode (version M16, Ensembl 91) as reference. We used mapped reads counts to call differentially expressed genes using DESeq2 (version 1.30.0) (Love *et al*, 2014), using the following setup as a model: *y* ∼ *stimulation*, where *y* are the normalized RNA gene expression levels and *stimulation* is either BDNF (1, 6, 10h) KCl (1, 6, 10h) or control (all samples). PCA analysis indicates that control neurons cluster together regardless of time point. For this reason, we compared library size normalized gene expression levels of BDNF, KCl in any time point versus all control samples, estimating in each case log2 fold changes, standard errors and significance using the Wald test implementation (two-sided). Genes with adjusted *P*-value < 0.1 were defined as DE-genes. To study variability in gene expression using unsupervised clustering, DE-genes are selected by scaled normalized gene expression variability across all samples. We clustered the mean-corrected expression changes in each gene using partitioning around medoids (PAM) clustering, setting the number of *k* medoids as 10. We compared the enrichment of gene ontology terms in each cluster versus other clusters using topGO (Alexa *et al*, 2006) version 2.42.0, using all mapped genes as genome background.

### ATAC-seq sample preparation and sequencing

For ATAC-seq, 50,000 mouse primary cortical neurons and 100,000 human iNeurons were harvested by centrifugation at 500 g x 5 min at 4 oC. Cell pellets were washed with an ice-cold PBS buffer and 50 mL cold lysis buffer (10 mM Tris-HCl pH 7.4, 10 mM NaCl, 3 mM MgCl_2_, 0.1% NP-40). The cell suspension was centrifuged at 500 *g* for 10 min at 4^0^C and supernatant was discarded. The nuclei-enriched pellet was immediately used for transposition reaction using Nextera DNA Library Prep Kit (Illumina, FC-121-1030). The transposition reaction was incubated at 37^0^C for 30 min and followed immediately by purification using QIAGEN MinElute Kit (Qiagen, 28204). Following steps were primarily based on the protocol described by Buenrostro *et al*., 2013 with minor changes. Purified DNA was subjected to an initial step of PCR amplification consisting of 5 cycles using NEBNext High-Fidelity 2X PCR Master Mix (NEB, M0541S) and standard barcoded primers of Nextera kit for each sample. We used 5 μl of partially-amplified libraries from each sample to perform quantitative PCR (qPCR) to determine the additional cycles for PCR amplification that were required while avoiding the saturation of the PCR, which may reduce the complexity of the original libraries. This was measured by comparing cycle numbers against 1/3 of the maximum fluorescence intensity using real-time PCR graphs on ABI7900 (Applied Biosystems). After the additional PCR cycles, samples were purified using Agencourt AMPure XP magnetic beads (Beckman Coulter, A63880). The quality of the DNA libraries was tested with the High Sensitivity DNA Bioanalysis Kit (Agilent, catalog # 5067-4626). DNA concentration of each library was measured with the Qubit dsDNA HS Assay Kit & fluorometer (ThermoFisher, Q32851). Based on the concentrations of each library, 8 samples were pooled together with each sample double barcoded for pair-end sequencing on NextSeq 500 platform EMBL, Heidelberg Gene Core facility.

### Chromatin accessibility computational data processing

We mapped ATAC-seq reads in each sample to the *M. musculus* genome (build mm10) using bowtie2 (v2.3.4.1) (Langmead & Salzberg, 2012). Mapped reads in each sample were used to call peaks in each treatment and time point with MACS2 (v2.2.7.1) (Zhang *et al*, 2008), using the following parameters to call peaks in each paired-end sample: “*--nomodel --shift -75 -- extsize 150*”. Then, we jointly analyzed called peaks and ATAC-seq reads to call differentially accessible peaks using DiffBind **[Stark R, Brown G (2011). DiffBind: differential binding analysis of ChIP-Seq peak data]**. We only considered peaks detected in at least two samples. We generated consensus peaks using an overlap of 66%. To correct counts per peak in all ATAC-seq datasets we used a LOESS normalization step. Normalized ATAC-seq peaks counts were obtained with DESeq2 using the following model: *y* ∼ *stimulation*, where stimulation is a treatment and time combination (e.g. BDNF_1h, KCl_6h). Similar to RNA-seq analysis, based on PCA grouping of control samples we decided to pooled all control samples and used them as a reference for comparison. Log2 fold changes between conditions and controls were assessed using Wald test (two-sided), defining DA-peaks when adjusted *P* < 0.1. Gained or closing DA-peaks are based on positive or negative log2 fold changes versus the normalized means of control samples, respectively.

### Chromatin accessibility epigenomics and gene ontology enrichments

General genomic annotations for gained and closing DA-peaks in each treatment-time point pair were done using HOMER (Heinz *et al*, 2010). To assess the enrichment of neuronal-specific chromatin marks, we used a previously reported hidden markov model generated from multiple chromatin marks and ChIP-seq data for neurons, using ChromHMM (Ernst & Kellis, 2012). As this model uses 15 states generated using the mouse genome build mm9, to interrogate our DA-peaks we converted these annotation ranges to mm10 using `liftOver` with default parameters. We report the log2 fold enrichment between the number of nucleotides in one of the 15 states, versus the number of nucleotides overlapping other states, using the function `OverlapEnrichment` of ChromHMM . To assess Gene Ontology enrichment of peaks proximal to DA-peaks we used a binomial and hypergeometric tests as implemented in the GREAT server (McLean *et al*, 2010), with default parameters to map peaks to genes: Peaks are associated to genes if located upstream of a Transcription Start Site (TSS) up to 5000 bp, downstream of a TSS up to 1000 bp, or up to 1 Mbp away from a TSS. We used as background regions unchanged peaks and their proximal target genes.

### Genomic data co-variation and loop data analysis

We defined Distal Regulatory Elements (DREs) as regions less than 2kbp and no further than 50Kbp from TSS regions. We assessed the enrichment of associations between DREs and DE-genes by sign quadrants, counting associations between peaks and genes. When two peaks are linked to one gene, to avoid duplicate counts we considered the sign of the peak whose change had the lowest adjusted *P* value. Enrichment was done using 2 x 2 contingency tables between double-positive and double-negative quadrant counts, using Fisher’s exact test. To assess the enrichment in the number of observed DA-peaks and DE-genes linked to each other, we compared the observed associations versus the ones defined through a permutation approach where log2 fold changes are maintained and distance values are scrambled across DA-peaks and DE-genes. This allows maintaining the structure of accessibility changes and number of significant events time points, while assessing the over-representation of genomically closing peaks and genes. We repeated this permutation approach 1,000 times and calculated Z-score for peaks (Z-peaks) and genes (Z-genes). An over-representation of associations between DA-peaks and DE-genes we defined when the Z-scores for both peaks and genes were greater than 2.5.

Similar to DE-genes, DA-peaks were also assessed by clusters of variability using PAM clustering using *k* = 10. DA-peak clusters were annotated by genomic location using HOMER (TSS or others). To assess the over-representation of DE-genes going up or down in each cluster, we calculated 2 x 2 contingency tables for closest DE-genes up in each cluster versus down in other clusters, or down in cluster versus up in other clusters. We assessed the significance in both cases with Fisher’s exact test, with BH correction.

ATAC-seq peak pair correlations were generated using all of the peak-pairs closer than 50 Kbp. ATAC peaks were stratified by none, one or both peaks annotated as DA-peaks in at least one contrast versus control samples. To compare these significance stratified regions with Hi-C contacts (Bonev *et al*, 2017), we allow a distance threshold of 10Kbp between Hi-C peak anchors and each ATAC-seq peak. Correlations were stratified by cell type using annotations by Bonev *et al*., based on shaman scores. This indicates higher covariation between peaks with known chromatin contact information.

Enrichment of same direction DA-peaks and DE-genes for Hi-C regions was done as follows: For each distal DA-peak, a Hi-C score was calculated versus the TSS region of target genes. If the shaman score was positive, then the association was considered strong. Counts for strong versus non-strong pairs were calculated for peak and gene associations both positive or negative, versus opposite sign directions, and assessed for significance using Fisher’s exact test (**Supp Fig. 4d**). Multiple testing correction was done using Benjamini Hochberg’s procedure.

### Motif enrichment analysis

We defined summit centered 200-bp regions from all differentially accessible peaks as foreground regions and retrieved background regions for each one using GENRE (Mariani *et al*, 2017), using a custom mm10 background. Briefly, a representative background sequence set is generated from a mouse specific database of reference genomic regions, matching for equivalent GC-content and CpG frequency, promoter overlap (extent of the sequence located within 2 kb upstream of a TSS), and repeat overlap. We mapped Transcription Factor (TF) motifs used in foreground and background sequences using *(i)* a *8*-mers reference set for 108 TF specificity groups generated from Protein Binding Microarray (PBM) data (Mariani *et al*, 2017), and *(ii)* a extensive database of Position Weight Matrices (*PWM*s) retrieved from the CIS-BP database (Weirauch *et al*, 2013). For mapping of *8*-mers, we used GENRE to define a single score per sequence as the best E-score greater than 0.35, or -1 otherwise, using the PWMmatch function (R Biostrings) if the TF is represented as a PWM, and *grepl* if represented as a *k*-mer. To scan PWMs from the CIS-BP database, we used FIMO (Grant *et al*, 2011) with default parameters, and kept the highest score for downstream analyses. We used these scores to assess sensitivity and specificity using a Receiver Operating Characteristic (ROC) analysis in the classification of foreground versus background sequences as in each treatment and time point, respectively. We used the Area Under the Curve (ROC-AUC) to define significantly enriched TF specificity modules and motifs. To assess significantly enriched modules, we used Wilcoxon rank sums tests, one-sided, adjusted with a Benjamini Hochberg procedure (FDR = 10%). If significant, we used a ROC-AUC threshold of 0.55 for interpretation of enriched TFs. To show TF expression levels, we used both the TF family annotations provided by CIS-BP and the TF specificity annotations from GENRE.

### TF motif co-enrichments in DA-peaks

Top five significant motifs by highest ROC-AUC values observed were assessed for co-enrichment in DA-peaks. Fold Enrichment and significance of group-wise over-representations were assessed using the hypergeometric test, implemented in the package SuperExactTest (Wang *et al*, 2015).

### Associations between bZIP and co-regulatory TFs with accessibility and expression changes

We assessed the association of presence of bZIP and co-factors motifs to changes in accessibilities and gene expression. Briefly, we compared log2 fold changes in the chromatin accessibility values for peaks with only bZIP motifs or bZIP and any co-regulatory motif. This analysis was also repeated for the expression values of closest genes to mapped peaks, and for genes with mapped peaks harboring in TSS regions. Statistical differences were assessed using a Wilcoxon rank sums test, two-sided, and adjusted using Benjamini Hochberg’s procedure.

### CTCF specific analyses at differentially accessible peaks

To assess the enrichment of DA-peaks for CTCF promoter-exon loops we used a previously released dataset of promoter-exon contacts (Ruiz-Velasco *et al*, 2017) to compare the odds ratio between DA-peaks in these loops versus unchanged peaks. We used Fisher’s exact test to assess the overrepresentation of DA-peaks in those regions, versus unchanged peaks in those regions. Additionally, we assessed the enrichment of a convergent versus divergent CTCF motif orientations across all proximal ATAC peak-pairs (less than 50Kbp) and regardless of promoter-exon loops, using the CTCF PWM for this (CIS-BP ID: M06483_1.94d) (Weirauch *et al*, 2013). Enrichment of DA-peaks in exons and introns was done using 2 x 2 contingency tables by time point (1, 6, or 10h) and DA-peak label (gained, closing or unchanged). Adjusted *P* values in these analyses were corrected using Benjamini Hochberg’s procedure.

### Differentially used exons computational and validation analyses

We called differentially used exons (DUEs) using mapped RNA counts across all samples in time point 1 hour (Gencode version M16). We used genes with non-zero counts in at least one treatment and time point to perform comparisons between BDNF and KCl in matched timepoints, using DEXSeq (Anders *et al*, 2012). *P-*values are adjusted using Benjamini Hochberg’s procedure (FDR less than 10%).

DUE for validation were based on the presence of a promoter-exon CTCF loop reported by Ruiz-Velasco *et al*. (Ruiz-Velasco et al, 2017). Primary cortical neuron cultures on DIV 7 were stimulated with BDNF or KCl for 1 hour, each condition performed on independent biological duplicates, and RNA was isolated for expression analysis of the DUEs via RT-qPCR. Primers for exons of three gene examples were selected, in addition to an additional same-gene exon for internal control.

For Trio, exon 29 with additional 3’UTR (E29+3’UTR) sequence is a DUE between BDNF and KCl in time point 1 hour. Three primer sets were designed to differentiate E29 (+3’UTR) from constitutive exon 28 (E28). In two of the sets, the forward primer lies within the coding region of E29 with a reverse primer in the immediately downstream 3’UTR region, whereas the third set did not include the 3’UTR region. For the constitutive exon, both forward and reverse primers lie in the coding region of E28. For Stxbp5 primer sets were designed to allow selective amplification of E01 against a constitutive exon, E05. For Cpe-201 primer sets were designed, one for E06 which is a DUE and another for constitutive E09. All primer sequences used to perform qPCR are as follows: Trio E29+3’UTR Forward set 1 (5’ CTCAGAGCAACGGGGTAAGAG 3’), Reverse set 1 (5’ GTGCTGGAGAGCTGGAGTTAG 3’), Forward set 2 (5’ GGCACCTTGACACCCACCT 3’) and Reverse set 2 (5’ GACCACAAAATGAGCCGGGA 3’); Trio E29 (without 3’UTR region) Forward (5’ CAAGCTCCTTCACCTTCCCG 3’) and Reverse (5’ CCTGGCACTCTTACCCCGTT 3’) ; Trio E28 Forward (5’ TGAGTTGCCTCTGCTTGGAG 3’) and Reverse (5’ GGACGCTTGGACTGGATGAA 3’); Stxbp5 E01 Forward (5’ CAACATCAGGAAGGTGCTGG 3’) and Reverse (5’ GAAGTGTTCGGACTGGAGCG 3’); Stxbp5 E05 Forward (5’ TGCCATCTGCCTTTCCAGAG 3’) and Reverse (5’ TGACATAGCCTGAGAGTGTGA 3’); Cpe-201 E06 Forward (5’ TGCTTCGAGATCACTGTGGAG 3’) and Reverse (5’ CTGCTCCAGGTAGCTGATGA 3’); and Cpe-201 E09 Forward (5’ TGTCTGGATCTACTTCATTCTTACA 3’) and Reverse (5’ CGCAGTACAGGGTTCACAGA 3’).

The stimulation-dependent fold change of each exon was quantified from 3 technical qPCR replicates normalized against Rpl13 as a reference gene. Fold changes relative to Rpl13 values were used to calculate the exon ratio between each tested exon and their internal exon as a reference. Comparisons between groups were assessed for statistical significance using *t*-tests (two-sided).

### Subcellular proteome analysis by mass spectrometry

Stimulated mESC-derived neurons were harvested using a scraper and pelleted at 500 *g* for 5 min. Cell pellets corresponding to 10 million cells were washed twice with ice-cold PBS and subjected to subcellular protein extractions using a Subcellular Protein Fractionation for Cultured Cells kit (Thermo Fisher Scientific, TFS #78840) following manufacturer’s instructions. Each condition (Ctrl, BDNF and KCl stimulations) were assayed in duplicates.

From each subcellular fraction, 10 ug of denatured protein, free of nucleic acids, were subjected to sample preparation for MS using a modified SP3 method (https://doi.org/10.15252/msb.20145625). Briefly, protein samples were precipitated onto Sera-Mag SpeevBeads (GE Healthcare, #45152105050250 and #65152105050250) using filter plates (Millipore #MSGVN22) for acidification and washes. Proteins were digested with trypsin and Lys-C, and the resulting peptides were vacuum dried and labeled with TMT labels (TMT10plex and TMT11, TFS, #90110, #A37724). For each replicate, labelled peptides from the following fractions were pooled to form a TMT11 set: unstimulated, BDNF-stimulated and KCL-stimulated, cytosolic (CEF), nuclear (NE), and chromatin-bound (CHR) fractions (channels 1-9), as well as a membrane fraction (channel 10) and a cytoskeletal fraction (channel 11) pooled from all three different conditions to increase coverage. Pooled peptides were desalted, washed and vacuum dried on OASIS HLB plates (Waters 186001828BA). Before LC/MS analysis, peptides were pre-fractionated into 12 fractions by Ultimate 3000 (Dionex) HPLC high-pH reverse chromatography and vacuum dried.

Reconstituted peptides were analysed by nanoLC-MS/MS on an Ultimate 3000 RSLC (TFS) connected to a Q Exactive Plus (TFS) mass spectrometer. Peptides were loaded on a trapping cartridge (Acclaim C18 PepMap 100, TFS) using 0.1% FA (solvent A) and separated on an analytical column (nanoEase M/Z HSS C18 T3, Waters) with a constant flow of 0.3 µl/min applying a 120 min gradient of 2–40% of 0.1% FA in CAN (solvent B) in solvent A. Peptides were directly analyzed in positive ion mode. Full scan MS spectra with a mass range of 375– 1200 m/z were acquired in profile mode using a resolution of 70,000 (maximum fill time of 250 ms or a maximum of 3e6 ions (AGC)). Precursors were isolated using a Top10 method with an isolation window of 0.7 m/z, fragmented using 30 NCE (normalized collision energy), and MS/MS spectra were acquired in profile mode with a resolution of 35,000, and an AGC target of 2e5 with a dynamic exclusion window of 30 s.

### Proteomic computational data analysis

Mass spectrometry raw files were processed using IsobarQuant (https://pubmed.ncbi.nlm.nih.gov/26379230/) and peptide and protein identification was obtained with Mascot 2.5.1 (Matrix Science) using a reference mouse proteome (uniprot Proteome ID: UP000000589, downloaded 14.5.2016) modified to include known common contaminants and reversed protein sequences. Mascot search parameters were: trypsin; max. 2 missed cleavages; peptide tolerance 10 ppm; MS/MS tolerance 0.02 Da; fixed modifications: Carbamidomethyl (C), TMT16plex (K); variable modifications: Acetyl (Protein N-term), Oxidation (M), TMT16plex (N-term).

IsobarQuant output data was analyzed on a protein level in R (https://www.R-project.org) using an in-house data analysis pipeline. In brief, protein data was filtered to remove contaminants, proteins with less than 2 unique quantified peptide matches, as well as proteins, which were only detected in a single replicate. Subsequently, protein reporter signal sums were normalized within each subcellular fraction across the two TMT sets (replicates) and across the three conditions using the vsn package (https://pubmed.ncbi.nlm.nih.gov/12169536/). Significantly changing proteins between BDNF and KCL stimulation conditions in the CHR subcellular fraction were identified by applying a limma analysis (https://pubmed.ncbi.nlm.nih.gov/25605792/) on the vsn-corrected signal sums. Replicate information was included as a covariate to adjust for batch effects caused by the separate TMT labelings and MS runs. T-statistics and p-values were obtained using the eBayes function from the limma package, and resulting p-values were corrected for multiple testing using the Benjamini-Hochberg method with the topTable function in the limma package.

### Linkage disequilibrium score regression analysis

For each group of Chromatin accessible regions, we performed partitioning heritability analyses to assess the proportion of variants associated to human complex traits harboring or proximal to differential accessible peaks. For mouse neurons we used an adjusted P-adjusted threshold of 0.2 for peak selection. In the case of hIPSC-derived neurons, due to the low number of DA-peaks detected we used raw P-values for peak selection, and the same threshold. Annotations for mouse neurons and hIPSC-derived neuron samples were extended to 10 Kbp, allowing a recovery of local genetic variants in the context of peaks, and genetic signal of proximal linkage disequilibrium (LD)-associated regions to be recovered.

We collected GWAS studies from two sources (i) a dataset of neuronal complex trait cohort pre-processed using steps described by (Hook & McCallion, 2020), and (ii) neuronal and non-neuronal complex traits GWAS cohort data preprocessed by Dr. Alkes Price group and available at https://alkesgroup.broadinstitute.org/sumstats_formatted. A description of the datasets and main features is provided in **Supplementary Data 4**.

Mouse regions were converted to human regions using *liftOver* and for all consensus peaks. Conserved blocks with length longer than 10,000 bp are discarded, recovering a total of 16,580 peaks mapped in the human genome.

Using as a reference genetic population data from 1000 Genomes Europeans (Auton *et al*, 2015), genetic variants in each study were pre-processed to leverage LD-block information. This modeling allows aggregation of LD-genetic associated GWAS SNPs to be aggregated as tags and increasing the strength of association between genomic annotations and polygenic trait dataset. Regression scores for gained DA-peaks, closing DA-peaks, all DA-peaks (DA) and consensus chromatin peaks are independently interrogated against all GWAS studies by computing partitioned heritability coefficients, using the package *ldsc* (Finucane et al, 2015). Significance *P*-values associated with the estimate of the first regression coefficient are adjusted for multiple testing correction using Benjamini Hochberg’s procedure.

## Supplementary Data

**Supplementary Data 1.**

Log2 fold changes changes in mouse gene expression and chromatin accessibility, including statistics and adjusted p-values.

**Supplementary Data 2.**

Coordination between DA-peaks and proximal DE-genes observed between all treatment and time combinations, measured as Z-scores.

**Supplementary Data 3.**

Molecular Biology readouts based on qPCR for *Arc* gene expression, and exon ratios to measure differences in exon abundances for *Trio, Stxbp5* and *Cpe*.

**Supplementary Data 4.**

Description of GWAS summary statistics and LSDC coefficients for mouse DA-peaks.

